# High confidence glycosomal membrane protein inventory unveils trypanosomal Peroxin PEX15

**DOI:** 10.1101/2023.10.13.562043

**Authors:** Chethan K. Krishna, Hirak Das, Lisa Hohnen, Wolfgang Schliebs, Silke Oeljeklaus, Bettina Warscheid, Vishal C. Kalel, Ralf Erdmann

**Affiliations:** Department of Systems Biochemistry, Institute of Biochemistry and Pathobiochemistry, Faculty of Medicine, Ruhr University Bochum, Bochum, Germany; Faculty of Chemistry and Pharmacy, Biochemistry II, Theodor-Boveri-Institute, University of Würzburg, Würzburg, Germany

**Keywords:** peroxisome, glycosome, cell fractionation, mass spectrometry, peroxin, peroxisomal membrane protein, PEX15

## Abstract

Infections by trypanosomatid parasites cause Chagas disease, Human African Trypanosomiasis, and Leishmaniasis, affecting over 12 million people worldwide. Glycosomes, the unique peroxisome-related organelles of trypanosomes are essential for their survival, and hence their metabolic functions and biogenesis mediated by peroxins (PEX) are suitable as drug targets. Here we report on a comprehensive protein inventory of glycosomal membranes through advanced subcellular membrane protein profiling employing quantitative mass spectrometry. Our quantitative analysis resulted in the identification of 28 novel high confidence glycosomal membrane proteins. Our in-depth protein inventory of glycosomal membranes serves as an important resource for characterizing glycosome biology and drug development. We validated four so far unknown glycosomal membrane proteins, including two tail-anchored (TA) proteins, a homolog of human peroxisomal PXMP4, and a Macrodomain-containing protein. Using a structure-based approach, we identified one of the TA proteins as the long-sought *Trypanosoma* PEX15. Despite its low sequence similarity, *Trypanosoma* PEX15 exhibits structural and topological similarities with its yeast (Pex15) and human counterparts (PEX26). We show that PEX15 is an essential integral glycosomal membrane protein that interacts with PEX6. Accordingly, RNAi knockdown of PEX15 in bloodstream form trypanosomes demonstrates that it is essential for glycosome biogenesis and parasite survival. Considering the low degree of conservation with its human counterpart, PEX15 is a promising molecular target for drug development.

## Introduction

Kinetoplastids are a group of flagellated protists with a DNA-containing region called kinetoplast found within their single mitochondrion [1]. Trypanosomatids are a subgroup of kinetoplastids that share the features of having a kinetoplast and a flagellum [2]. The insect-borne infections of trypanosomatid parasites cause different human diseases. These include Chagas disease, Human African Trypanosomiasis (HAT), and Leishmaniasis, which are caused by *Trypanosoma cruzi*, *T. brucei*, and *Leishmania* species, respectively, posing a major concern for public health. The incidence of trypanosomiasis affects approximately 6-7 million people and a population of around 75 million at risk [3, 4]. Leishmaniasis (all forms of diseases) exhibits a prevalence of over 6 million cases and over 600 million people at risk, with around 1 million new cases of cutaneous leishmaniasis and 50,000-90,000 cases of visceral leishmaniasis annually [3–6]. Presently, treatment options are available for the early stages of HAT [7] and, to some extent for visceral leishmaniasis, with candidates in clinical trials [8]. However, treatments for Chagas disease and other forms of leishmaniasis still face challenges due to significant side effects and toxicity [9]. There is a requirement to identify novel drug targets and develop new therapies to counter these infections.

Trypanosomatid parasites have specialized peroxisomes called glycosomes that compartmentalize glycolytic enzymes [10, 11]. This compartmentation is unique because, in other organisms, glycolysis occurs completely in the cytosol. The compartmentation is crucial for parasite survival because the mislocalization of glycolytic enzymes, which lack feedback inhibition, is lethal for the parasites [12, 13]. Glycosomes also contain enzymes for other pathways like gluconeogenesis [14], pentose phosphate pathway, pyrimidine biosynthesis, and purine salvage pathways [15]. Depending on their life cycle stage, glycosomes also contain enzymes for ether-lipid biosynthesis [16]. The organelle’s various functions make it essential for the different life cycle stages of the parasites. In this aspect, biogenesis of glycosomes has emerged as an attractive drug target. RNA interference-mediated knockdown of expression of various peroxins is lethal for the parasites (reviewed in [17]). Furthermore, small molecule inhibitors that specifically disrupt glycosome biogenesis have been developed which show therapeutic effects *in vivo* [18].

Like peroxisomes, glycosomes do not possess DNA, and hence their proteins are synthesized in the cytosol, and imported post-translationally into the organelles [19, 20]. The biogenesis of these organelles depends on proteins collectively referred to as peroxins. The peroxisome/glycosome biogenesis depends on two distinct machineries for the targeting of proteins. One machinery is responsible for peroxisomal matrix protein import, while the other machinery is responsible for formation of the peroxisomal membrane [21]. Peroxisomal/Glycosomal membrane proteins (PMPs) contain membrane peroxisome targeting signals (mPTS) and depend on the cytosolic receptor PEX19 and membrane docking factor PEX3 and/or PEX16 for their targeting. Glycosomal matrix proteins contain peroxisomal targeting signals at the extreme C-terminus (PTS1) or close to the N-terminus (PTS2) [22, 23]. PEX5 is the cytosolic receptor for PTS1-containing proteins and a co-receptor of PEX7, the receptor for PTS2-containing proteins [24]. Following cargo release, the monoubiquitinated receptor is recycled back to the cytosol by the AAA-ATPase complex PEX1-PEX6 in an ATP-dependent manner [25, 26]. This complex along with the RING finger peroxins (PEX2, 10 and 12) form the exportomer, which functions in receptor ubiquitination and recycling [27–29]. In the cytosol, the receptor is made available for the next round of import, indicating that the receptor recycling is crucial for biogenesis of peroxisomes. In the absence of recycling, PEX5 accumulates at the membrane, leading to pexophagy in yeast [30]. Both AAA+ ATPases PEX1 and PEX6 have been identified in trypanosomes [31, 32]. However, the protein anchoring the PEX1–PEX6 complex, the suspected PEX15/PEX26 at the glycosomal membrane, remains unknown. Most PEX protein orthologs in trypanosomes were discovered by their primary sequence similarity with the known PEX proteins in other organisms, except PEX3 and the so far unknown PEX15/PEX26. Trypanosomatid PEX3 could be identified through structural homology search using HHPRED or Phyre2 [33, 34], however, all bioinformatics approaches failed to identify trypanosomal PEX15.

In recent years, subcellular proteomics approaches have been applied to uncover the composition of various organelles, particularly mitochondria, where databases like MitoCoP (Mitochondrial high Confidence Proteome) and MitoCarta3.0 have been established [35, 36]. Peroxisomal proteomics studies of different cell types and organisms have also been published [37–42]. Recently, Yifrach et al. performed a systematic multi-level analysis of the yeast peroxisomal proteome termed Peroxi-ome, expanding the peroxisomal proteome by 40% and revealed new peroxisomal functions [42]. Peroxisomes are versatile organelles that can rapidly adapt their proteome. Similarly, the composition of glycosomes also changes as the parasites cycle between insect and mammalian hosts upon infection and undergo various stages of differentiation. The spatial proteome of insect and host life cycle stages of African trypanosomes [43] and *L. donovani* has been described [44]. These global sub-cellular localization studies report lists of glycosomal proteins, but they were not designed to distinguish between membrane and matrix proteins. Several studies have also reported on the glycosomal proteome of *T. brucei*, *T. cruzi,* and *Leishmania* species [44–47]. However, the current glycosomal proteome is dominated by highly abundant glycosomal matrix proteins, particularly enzymes of the glycolytic pathway.

Eleven of the 16 known parasite peroxins are PMPs [13, 15]. Knowledge of all the components of the glycosomal protein import machinery (peroxins) will be clinically significant, particularly as novel potential drug targets. By interfering with these peroxins, the import of matrix or membrane proteins and, in turn, the biogenesis of glycosomes will be blocked. This will disrupt various glycosome-associated metabolic pathways, resulting in parasite death [48, 49]. In this aspect, the so far unknown PEX15/PEX26 of trypanosomes could be an ideal drug target owing to its very low expected sequence similarity with its human counterpart. Given the significance of PMPs as described above, a membrane protein inventory of glycosomes will provide insights into their functional importance, translocation mechanisms, and role in organelle biogenesis.

In this study, we describe the isolation of glycosomes and the enrichment of glycosomal membranes followed by quantitative mass spectrometry (MS) for protein correlation profiling (PCP) [41, 50, 51] to establish a high-confidence protein inventory of glycosomal membranes of *Trypanosoma brucei* (procyclic form). One of the major goals for establishing this inventory was towards identifying missing trypanosomal peroxins PEX15. Our approach led to the identification of 28 novel potential PMPs, of which 11 are likely exclusive glycosome localized, while the remaining 17 are dual or multiple localized membrane proteins. These proteins include novel potential peroxins, enzymes, metabolite transporters, proteins which might be involved in membrane contact sites, protein quality control, organelle transport or with unknown functions. We validated the subcellular localization of four newly identified glycosomal membrane proteins and we demonstrate that one of the identified PMPs is the so far unknown trypanosomal PEX15, a peroxin essential for glycosome biogenesis and parasite survival. *Tb*PEX15 and other identified trypanosomatid-specific PMPs, which do not have identifiable human counterparts, facilitate their prioritization as potential drug targets.

## Results

### Purification of glycosomes and enrichment of glycosomal membrane proteins

We employed a systematic approach to define the glycosomal membrane proteome, with the specific goal to identify low-abundant PMPs, including the elusive PEX15, that may function as parasite-specific peroxins, metabolite transporters, or proteins involved in membrane contact sites. To this end, we purified glycosomes from procyclic *T. brucei* cells based on differential centrifugation and density gradient centrifugation of an organelle-enriched fraction (**Fig. 1a**; [52]). Glycosomal membrane proteins were further enriched from glycosomal peak fractions of the density gradient by successive treatment with low-salt and high pH carbonate buffer (**Fig. S1;** [52]). Immunoblot analysis of density gradient fractions revealed that marker proteins for both the glycosomal matrix (aldolase, hexokinase, GAPDH) and membrane (GIM5, PEX11) were strongly enriched in fractions 12 and 13, indicating the presence of intact glycosomes (**Fig. 1b**). Marker proteins for mitochondria (mtHsp70, matrix; VDAC, outer membrane), ER (BiP, lumen), and cytoskeleton (tubulin) were largely depleted but still detectable in low amounts, while the cytosolic enolase is absent from these glycosomal peak fractions.

**Fig. 1:**
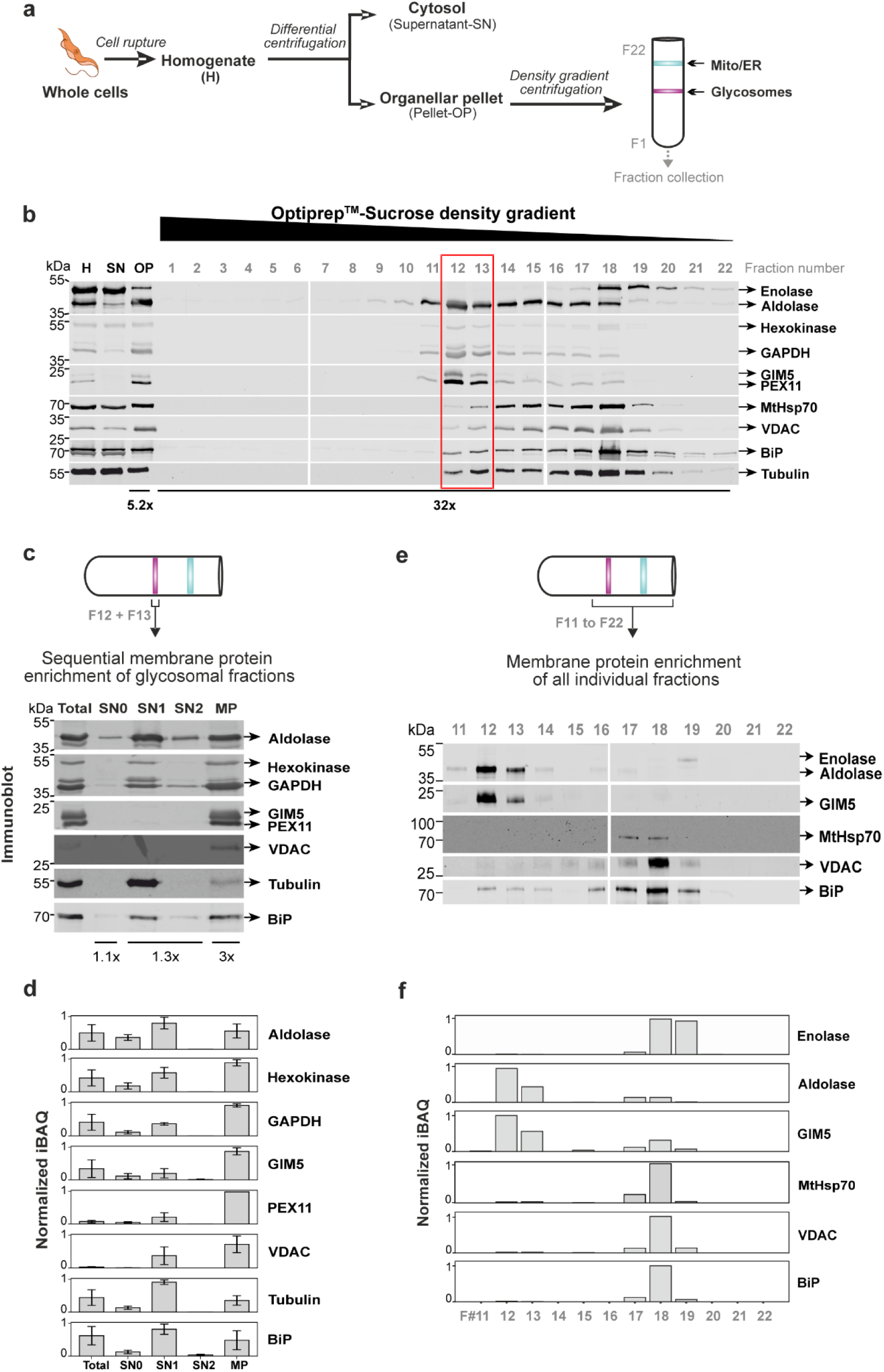
Preparation of a fraction enriched in glycosomal membrane proteins from *Trypanosoma*. **a.** Experimental workflow for the isolation of glycosomes from *T. brucei* cells using differential and density gradient centrifugation. The migration pattern of glycosomes (magenta) and other organelles (cyan) is shown schematically. F, fraction number. **b.** Fractions collected from the bottom of the density gradient were analyzed by immunoblotting using various antibodies as markers for subcellular compartments. The red box highlights the fractions enriched in glycosomes. H, homogenate; SN, supernatant representing cytosolic proteins; OP, organellar pellet. **c.** Glycosomes (F12 and F13 from **b**) were sequentially treated with low-salt buffer and twice with alkaline carbonate buffer to remove matrix proteins and peripheral membrane proteins. After each centrifugation step, supernatants and the final pellets were collected (protocol outlined in **Fig. S1**) and analyzed by immunoblotting. Total, intact glycosomes; SN0, supernatant of post low-salt treatment; SN1/SN2, supernatants post first or second carbonate treatment; MP, integral membrane-protein pellet. **d.** Samples obtained as described in (**c**) were analyzed by quantitative liquid chromatography-mass spectrometry. Shown is the mean (n = 3) of normalized MS intensities for the indicated proteins. Error bars indicate standard error of the mean. iBAQ, intensity-based absolute quantification. **e.** Immunoblot analysis of the carbonate-resistant protein pellet of gradient fractions 9-22. **f.** Abundance profiles of selected organellar marker proteins across density gradient fractions 11-22 based on normalized MS intensities (n = 1). Organellar marker proteins used in (**c-f**) are enolase (cytosol), aldolase, hexokinase, GAPDH (glycosomal matrix), GIM5, PEX11 (glycosomal membrane), mtHsp70 (mitochondrial matrix), VDAC (mitochondrial membrane), BiP (ER), and tubulin (cytoskeleton). Concentration factors relative to H or Total glycosomes are shown at the bottom in (**b**) and (**c**).

Glycosomal peak fractions (i.e., fractions 12 and 13) were pooled and further processed for the enrichment of glycosomal membrane proteins [53, 54]. To this end, glycosomes were sequentially treated with low-salt buffer and twice with alkaline carbonate buffer (pH 11.5) as outlined in **Fig. S1**. This resulted in the removal of glycosomal matrix proteins, proteins loosely associated with the membrane and any potential peripheral membrane proteins, to a certain extent. The final carbonate-resistant pellet (in the following referred to as membrane fraction) contained integral membrane proteins, which was confirmed by immunoblot analysis showing complete recovery of glycosomal membrane markers GIM5 and PEX11, as well as VDAC, a mitochondrial outer membrane protein (**Fig. 1c**). Surprisingly, portions of glycolytic enzymes (i.e., aldolase, hexokinase, GAPDH) of the glycosomal matrix remained in the membrane fraction, while tubulin was removed in the first carbonate extraction step. Peroxisomal and glycosomal enzymes are known to form crystalline cores [16, 55], which may remain resistant to the carbonate extraction. This is in line with observations that some glycosomal enzymes of *L. tarentolae* are urea-resistant [46]. The samples of the individual steps of the glycosomal membrane protein enrichment experiments were further analyzed by liquid chromatography-mass spectrometry (LC-MS), confirming the data of the immunoblot analysis and additionally providing a quantitative dimension to the results (**Fig. 1d, File S1**). The MS data clearly show that membrane proteins (i.e., GIM5 and VDAC) are largely enriched in the membrane fraction compared to the pooled glycosomal peak fractions of the density gradient (i.e., “Total” in **Fig. 1c**).

Furthermore, we prepared membrane protein-enriched fractions of individual density gradient fractions (i.e., fractions 11 – 22; chosen based on the protein distribution profile shown in **Fig. 1b**) and analyzed them by immunoblotting and LC-MS (**Figs. 1e, fand File S1**), which demonstrated that glycosomal membrane proteins (GIM5) clearly and distinctively peak in fractions 12 and 13, while mitochondrial and ER membrane proteins peak in fractions 18. Glycosomal matrix enzyme aldolase peaking in fractions 12 and 13 represents the carbonate resistant portion (as seen in **Fig. 1c**). We also observed a small portion of the cytosolic protein enolase in the membrane pellets of fractions 18 and 19. This is also observed for yeast enolase, which associates with mitochondria and remains resistant to carbonate treatment [56, 57]. Moreover, *Leishmania* enolase can also be found bound to membranes, specifically localized at the external surface of the plasma membrane [58].

### Defining the glycosomal membrane protein inventory

The strategy we employed to define a high-confidence inventory of the glycosomal membrane proteome of *T. brucei* relies on two different approaches employing quantitative MS to generate specific subcellular and suborganellar protein profiles (outlined in **Fig. 2a**) In the first approach, fractions obtained by sequential treatment of gradient-purified glycosomal peak fractions with low-salt and alkaline carbonate buffer to generate a fraction enriched in membrane proteins as described above (**Figs. 1b, c, d**) were profiled via label-free LC-MS. The resulting MS data were used to establish protein abundance profiles of well-defined sets of marker proteins for different subcellular (sub)compartments across the different membrane protein enrichment steps. Protein Correlation Profiling (PCP) [41, 50, 51] was then employed to determine the correlation between the distribution profile of individual proteins and the consensus profile of glycosomal membrane proteins, leading to the establishment of a preliminary list of putative glycosomal membrane proteins. Further classification provided a list of Class 1 and Class 2 candidates. In the second approach, glycosomal membrane protein candidates were determined based on their specific migration profiles measured across carbonate-resistant fractions of a density gradient using LC-MS (**Figs. 1e, f**). Prediction of transmembrane domains (TMDs), further literature and database curation and orthogonal validation of selected candidates resulted in the definition and characterization of a high confidence glycosomal membrane protein inventory (**Fig. 2a)**.

**Fig. 2:**
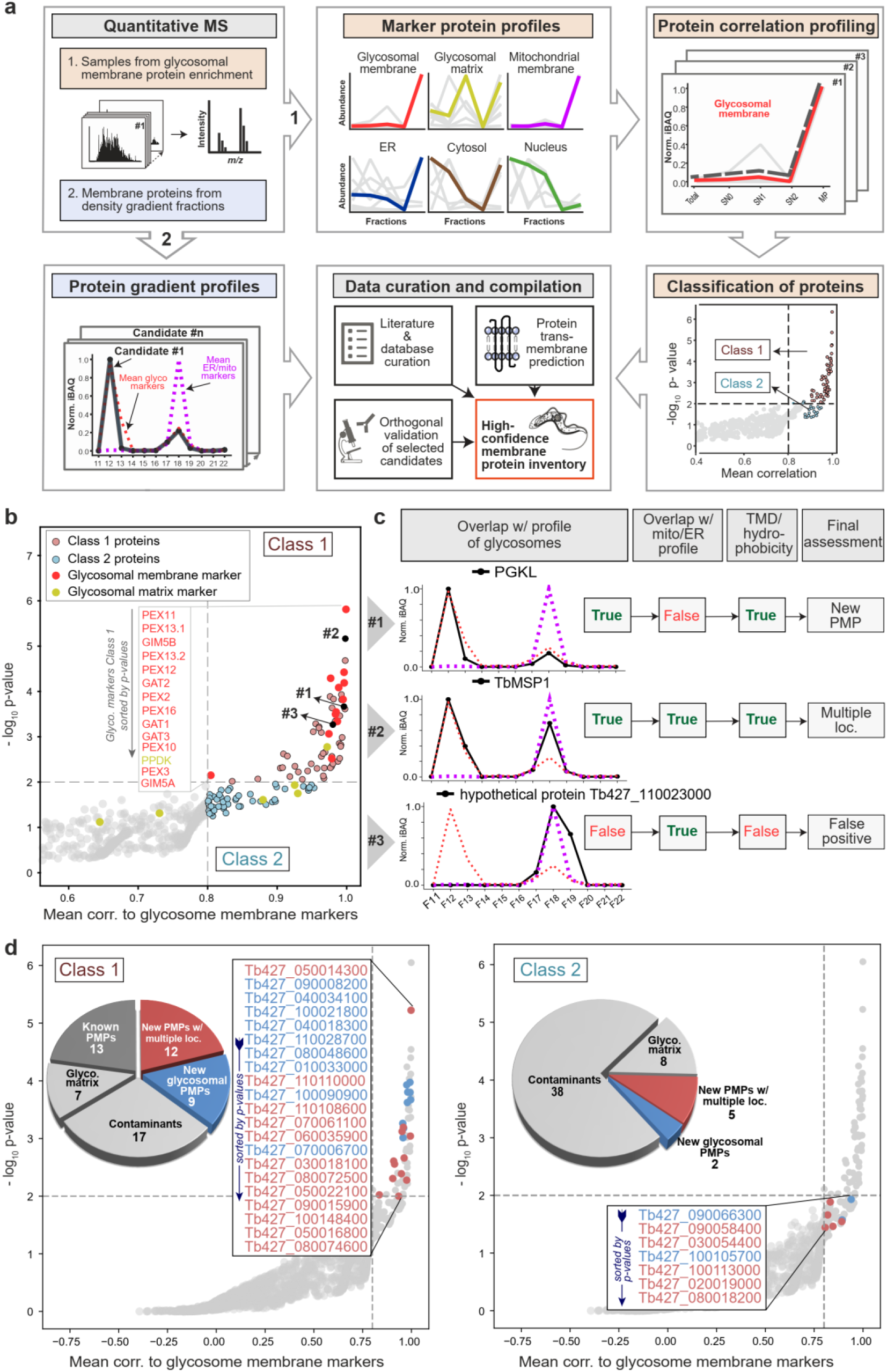
Establishment of a high confidence glycosomal membrane protein inventory using quantitative mass spectrometry. **a.** Overview of the strategy. The definition of the glycosomal membrane proteome was based on two quantitative MS datasets generated from (1) samples obtained in glycosomal membrane protein enrichment experiments (**Figs. 1c, d** and **Fig. S1**) and (2) carbonate-resistant membrane proteins of individual density gradient fractions (F11-F22; **Figs. 1e, f**). The first dataset was used to establish consensus profiles for selected organellar marker proteins across all samples of the carbonate extraction experiment. Protein correlation profiling was used to determine the correlation between the protein abundance profile of individual proteins and the consensus profile for glycosomal marker proteins. A preliminary list of glycosomal membrane proteins (Class 1 and Class 2) was defined based on the correlation and statistical significance (p-value) across the replicates (n = 3). The second dataset was used to verify the glycosomal localization of proteins and exclude contaminants based on protein abundance profiles across the density gradient. Prediction of transmembrane domains (TMDs), literature and database curation, and orthogonal validation of selected candidates completed the definition of the high confidence glycosomal membrane protein inventory. **b.** Rank product plot showing the mean correlation (corr.) between the abundance profile of individual proteins with the consensus profile of glycosomal membrane marker proteins across the samples of the glycosomal membrane protein enrichment experiments and the statistical significance (-log_10_ p-value) determined for proteins present in at least 2/3 replicates using the rank sum method. Candidates (mean correlation of ≥ 0.8) were divided into class 1 (p-value < 0.01) and class 2 (p-value of ≥ 0.01). **c.** Criteria applied to discriminate between new glycosomal membrane proteins predominantly localized in the glycosomal membrane (“New PMP”), proteins present in the membranes of glycosomes and other organelles (“Multiple loc.”), and false positives that were coenriched in carbonate-resistant membrane fractions. **d.** Composition of class 1 and class 2 proteins. Assignment of proteins to the individual subgroups is based on the abundance profile of individual class 1 and class 2 proteins in comparison to the consensus profile of glycosomal membrane and mitochondrial/ER marker proteins and the presence or absence of TMDs as indicated in (**c**). **a, b, d** Horizontal and vertical dashed lines indicate a p-value of 0.01 and a mean correlation of 80%, respectively.

Quantitative MS analysis of the samples obtained in glycosomal membrane protein enrichment steps (**Figs. 1c, S1**) led to the identification of 4,317 proteins (**File S1**). 2001 of these were (i) detected in at least two out of three biological replicates and (ii) present in at least two out of five samples per replicate and considered for further analysis by PCP. Based on the mean correlation between a protein’s distribution profile and the consensus profile of the glycosomal membrane marker proteins, a preliminary list comprising known and putative new glycosomal membrane proteins was established. We defined a mean correlation of ≥ 80% as threshold, since this cut-off encloses all glycosomal membrane markers (**Figs. 2b, S2a**), while the majority of glycosomal matrix marker proteins had a mean correlation of < 80% (**Fig. S2a**). This preliminary list was subdivided into PMPs of class 1 (p-value < 0.01; 59 proteins) and class 2 (p-value > 0.01; 53 proteins).

To assess the quality of our PCP-based approach, we analyzed the distribution of marker proteins for further subcellular localizations such as mitochondria, ER, nucleus and cytosol (**Fig. S2b).** None of the non-glycosomal marker proteins are among the class 1 candidate proteins, and in class 2, only the ER marker putative dolichol phosphate-mannose synthase is present, which underscores the potential of our approach to discriminate between glycosomal and non-glycosomal membrane proteins. Furthermore, we compared our results with data of a previously published quantitative proteomics study of affinity-purified glycosomes, which did not distinguish between matrix and membrane proteins [47]. Thirty of our class 1 proteins were reported to be present in the glycosomal proteome, while proteins categorized as “possibly glycosomal” were absent from our class 1 proteins (**Fig. S2c; File S1**).

The glycosomal peak fractions collected from the sucrose density gradients (**Figs. 1b, c, d**) still contained residual amounts of proteins from other organelles such as mitochondria and ER, which were coenriched in the carbonate-resistant membrane fractions. As a consequence, the consensus profiles of mitochondrial and ER proteins across the membrane protein enrichment steps resemble the consensus profile of glycosomal membrane proteins (**Fig. 2a**, top, second box). Thus, it can be expected that our preliminary list of class 1 and class 2 PMPs still includes non-glycosomal proteins. In addition, glycosomal membrane proteins may be present in further subcellular niches like the ER in case of *Tb*PEX13 [59], an information that is not clearly deducible from our PCP analysis but with possibly important implications for the functional significance of a protein. Thus, to identify glycosomal membrane proteins with multiple subcellular localizations, while removing contaminants from other organelles, we analyzed carbonate-resistant membrane pellets of density gradient fractions by quantitative MS (F11-F22; **Fig. 1f, File S1**). From this data, we established abundance profiles across the gradient for all preliminary class 1 and class 2 proteins identified in at least two fractions (**File S2**) and compared their profiles manually with the consensus profile of both glycosomal membrane and mitochondrial/ER marker proteins showing a peak in fraction F12 and F18, respectively (**Figs. 1e, f, 2c**). Furthermore, since some glycosomal matrix enzymes persisted carbonate extraction and, thus, were present in our preliminary list (**Fig. 2b**), we analyzed all candidate proteins for the presence of TMDs and/or hydrophobicity. This approach enabled us to distinguish between (1) glycosomal membrane proteins with main localization in glycosomes, showing exact overlap with the glycosomal profile and possessing known or clearly predicted TMDs or significant hydrophobic domains (**Fig. 2c**, example #1, putative phosphoglycerate kinase [PGKL]); (2) glycosomal membrane proteins with multiple localization, showing overlap with the profiles of both glycosomal and mitochondrial/ER marker proteins and TMDs/hydrophobicity (**Fig. 2c**, example #2, *Tb*Msp1); and (3) false positives, showing no overlap with the glycosomal profile and no TMDs/hydrophobicity (**Fig. 2c**, example #3, hypothetical protein Tb427_110023000).

After manual curation, the class 1 list consists of 34 proteins, including 13 known PMPs, 9 new glycosomal membrane proteins, and 12 new multi-localized PMPs. 24 contaminants were removed, of which 7 are glycosomal matrix proteins and 17 contaminants from other organelles (**Fig. 2d**, left). In addition, one protein (DEAD/DEAH box helicase, Tb427_100067300) was removed because it did not meet the criteria for establishing a profile across the density gradient fractions. By filtering of the list of class 2 proteins 45 contaminants (7 glycosomal matrix proteins, the peripheral glycosomal membrane protein PEX4, and 38 proteins from other organelles) were removed. Contaminant proteins originated mainly from mitochondria and the ER but also included ribosomal proteins. At the end, 7 novel glycosomal membrane proteins were identified in this list, including 2 that exclusively localize to glycosomes and 5 new multi-localized PMP candidates (**Fig. 2d**, right).

Interestingly, class 1 contains two proteins of the mitochondrial outer membrane (*Tb*Msp1 [Tb427_050014300] and a putative 3-oxo-5-alpha-steroid 4-dehydrogenase [Tb427_030018100]), as well as a mitochondrial inner membrane protein (Dienoyl Co-A Reductase [Tb427_030018100]) [60] (**Fig. S2d**). In fact, *Tb*Msp1, which shows a correlation of 99.8% with the consensus profile of the glycosomal membrane marker proteins (**Fig. 2b, File S1**), has recently been shown to exhibit a dual localization to glycosomes and the mitochondrial outer membrane [61]. Similarly, 3-oxo-5-alpha-steroid 4-dehydrogenase has been reported to be present in glycosomes as well [47]. Our analysis confirms the dual localization of both proteins based on their abundance profiles across the density gradient fractions (**File S2**).

The identified class 1 PMPs were further analyzed by bioinformatic analysis using sequence-homology (BLAST) or 3D structural homology using Foldseek [62]. This analysis led to the identification of four potential homologues of known peroxisomal membrane proteins.

The structure-based analysis identified the Transmembrane Protein 135 (TMEM135) and the Acyl/Alkyl DHAP Reductase (ADHAPR) in *T. brucei*. InterPro Scan analysis predicted that the putative *Tb*TMEM135 contains seven TMD’s and belongs to the TMEM135 family, while the putative short-chain dehydrogenase (*Tb*SDR) contains three TMD’s, is homologous to mammalian ADHAPR and is classified within the short-chain dehydrogenase family (**Figs. S3a, b**).

Additionally, two new putative members of the PEX11 family proteins in *T. brucei* were identified, based on their structural similarity to the known *Sc*PEX11 and *Tb*PEX11, as evidenced by superimposition with *Sc*PEX11 used as a reference **(Fig. S3c)**.

Taken together, multifaceted subcellular protein profiling provided a high confidence glycosomal membrane protein inventory with 28 new candidates for further exploration.

### From inventory to the validation: Characterization of shortlisted PMPs from the inventory

From the class 1 high-confidence glycosomal membrane protein list (**Table 1**), we shortlisted four candidates for a further characterization. For a diverse, representative set of candidates, selection was based on a mean correlation cutoff value of at least 0.95, with two candidates exhibiting highest and two average MS scores. The shortlisted proteins are peroxisomal membrane protein 4, macrodomain protein 2, and two hypothetical proteins tentatively named tail-anchored protein 1 (TA1) and tail-anchored protein 2 (TA2) (**Fig. 3a**). We particularly focused on the two TA proteins as they were promising candidates for the missing *Tb*PEX15 ortholog. The selected candidates were initially subjected to a sequence similarity search (BLAST) against yeast (*S. cerevisiae*), human, and plant (*A. thaliana*) protein databases, which identified an ortholog of the human PXMP4, to which we refer as *Trypanosoma* peroxisomal membrane protein 4 (*Tb*PMP4; Tb427_090008200). The other three proteins are only conserved in trypanosomatids and annotated as hypothetical proteins in the *Trypanosoma* database (TriTrypDB; www.tritrypdb.org). Bioinformatic analysis, such as TMD prediction using Phobius web tool and domain architecture prediction using InterPro Scan revealed that *Tb*PMP4 has four TMDs with a conserved Tim17 superfamily domain and a sequence identity of 36.9% with *Hs*PXMP4 (**Fig. S4a**). One of the three hypothetical proteins has a macrodomain with a single TMD close to the N-terminal region and will be referred to as Macrodomain 2 protein (MDo2). In a previous study, another macrodomain protein (MDo1) was identified and characterized in both *T. brucei* and *T. cruzi* [63]. Both *T. brucei* MDo1 and MDo2 contain a PTS1 signal characteristic of peroxisomal/glycosomal matrix proteins and are constituents of a high confidence glycosomal proteome [47]. Most interestingly, MDo2 contains a TMD in addition to its PTS1, a feature that previously only has been reported for PEX13.1 in *Trypanosoma* [64]. The remaining two hypothetical proteins, TA1 and TA2, harbor a single TMD near their C-terminus but lack any conserved domains. The domain architecture of all four shortlisted proteins is shown in **Fig. 3b**.

**Fig. 3:**
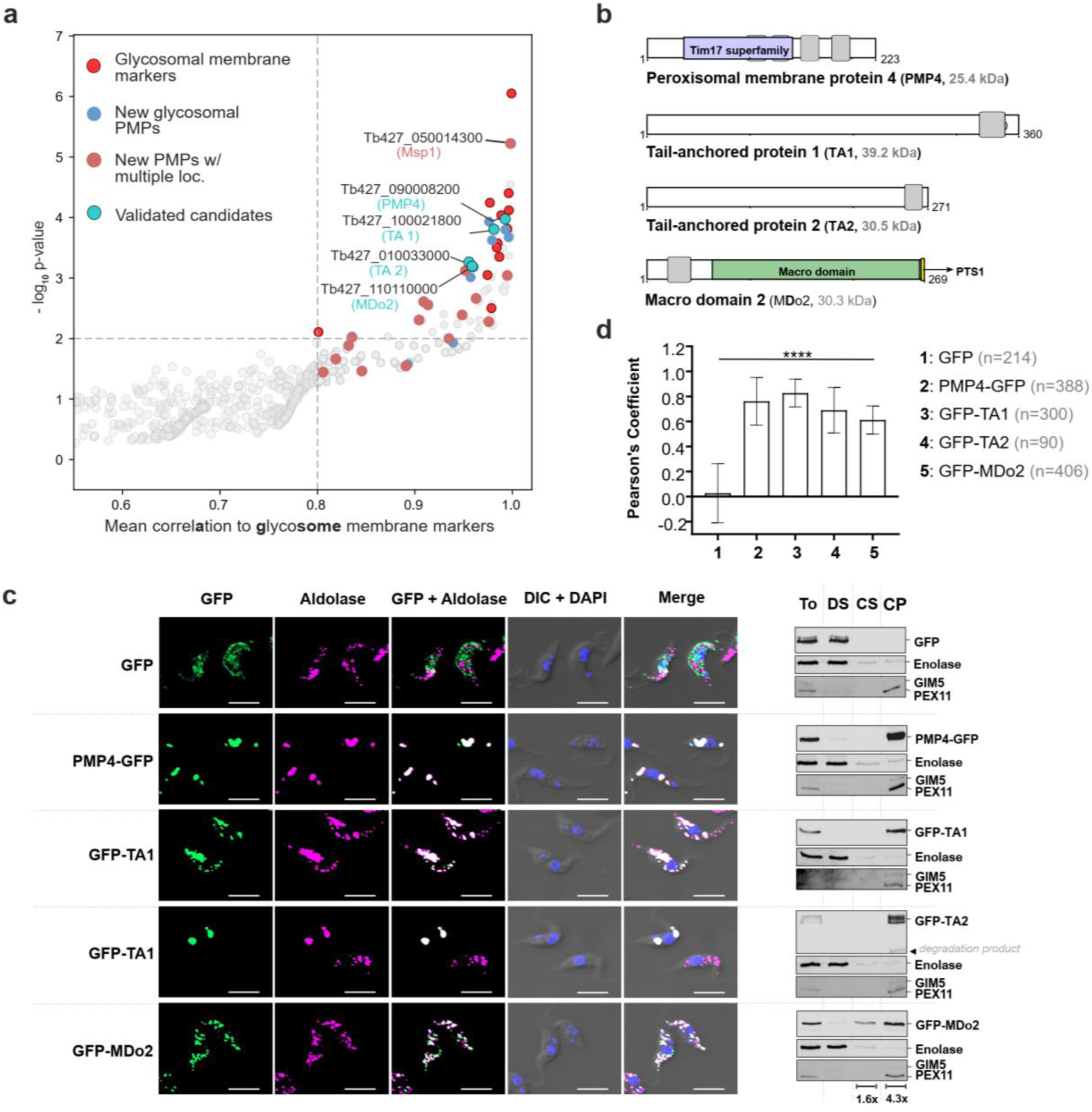
Validation of selected novel glycosomal membrane proteins. **a** Rank product plot as in Fig. 2b, highlighting known and newly identified putative PMPs after curation of candidates. Four candidate PMPs, which were further validated are marked in dark blue circles. **b** Domain architecture of the selected novel glycosomal PMP candidates: *Tb*PMP4 (Tim17 superfamily domain in blue), tail-anchored protein 1 (TA1), tail-anchored protein 2 (TA2) and *Tb*MDo2 (macrodomain in green, C-terminal PTS1 signal in yellow). Predicted transmembrane segments are depicted as grey boxes. **c** Immunofluorescence microscopy analysis of the indicated GFP-tagged PMP constructs demonstrating colocalization with the glycosomal marker aldolase (pseudocolored in magenta). DAPI stains nucleus and kinetoplast. The negative control GFP is cytosolic, as evident from the overall diffuse cell labelling. The scale bar represents 2 μm. Same cell lines were subjected to detergent extraction with digitonin, and alkaline carbonate, which demonstrates that the GFP-tagged proteins are integral membrane proteins. For the corresponding immunoblots, shown on the right, fractions of indicated cells were analyzed using anti-GFP antibodies to detect the fusion proteins along with control proteins enolase, GIM5 and PEX11, cytosolic and glycosomal membrane proteins, respectively. To: total cell lysate; DS: digitonin supernatant containing cytosolic proteins; CS: carbonate supernatant, containing organellar matrix and peripheral membrane proteins; and CP: carbonate pellet containing integral membrane proteins. Concentration factors relative to the total are depicted at the bottom. **d** Quantification of the colocalization of GFP-tagged PMPs to glycosomes. The Pearson’s coefficient of colocalization to respective organelle is shown. Error bars represent mean with standard deviations. Statistical significances were calculated by one-way ANOVA, with Dunnett’s multiple comparisons test (n ≥ 90 cells). ****, p < 0.0001.

**Table 1:** High confidence glycosomal membrane protein inventory: The table contains known PMPs (white) and newly identified potential PMPs (grey) in the order of their rank in our experimental analysis (>80% correlation in glycosomal PMP enrichment).

| Gene ID | Protein name | Description | TMD | Localization | Putative function |
| --- | --- | --- | --- | --- | --- |
| Tb927.11.11520 | PEX11 |  | 4 |  |  |
| Tb927.5.960 | Msp1 | Mitochondrial sorting protein 1 | 1 | Dual/multilocalized | Protein quality control |
| Tb927.9.9430 | PEX13.2 |  | 3 |  |  |
| Tb927.10.14720 | PEX13.1 |  | 2 |  |  |
| Tb927.11.2380 | PGK-Like | Phosphoglycerate kinase, putative | 4 | Glycosomal | Metabolism |
| Tb927.11.3130 | GAT2 |  | 6 |  |  |
| Tb927.4.1600 |  | Zeta toxin/AAA domain/Chromatin associated protein KTI12, putative | 1 | Glycosomal | ATPase |
| Tb927.9.1720 | PMP4 | Tim17/Tim22/Tim23/Pmp 24 family, putative | 4 | Glycosomal | Metabolite transporter |
| Tb927.10.15850 | PEX12 |  | 1 |  |  |
| Tb927.4.4050 | GAT1 |  | 6 |  |  |
| Tb927.3.2340 | PEX2 |  | 3 |  |  |
| Tb927.9.6450 | PEX16 |  | 2 |  |  |
| Tb927.11.1070 | GAT3 |  | 2 |  |  |
| Tb927.10.1860 | PEX15 |  | 1 (TA) | Glycosomal | Peroxin |
| Tb927.7.5540 | DECR | Dienoyl Co-A Reductase | 1 (TA) | Dual/multilocalized | Metabolism, C-terminal highly conserved YKI |
| Tb927.8.4370 |  | hypothetical protein, conserved | 4 | Glycosomal | Unknown |
| Tb927.11.10260 | PEX3 |  | 1* |  |  |
| Tb927.9.11600 | GIM5B |  | 1 |  |  |
| Tb927.4.3160 | DHAPAT /DAT | Acyltransferase, putative | 2 | Glycosomal | Metabolism (Ether lipid synthesis), highly conserved PTS1 SRM |
| Tb927.3.2410 | PEX10 |  | 2 |  |  |
| Tb927.6.3080 |  | hypothetical protein, conserved | 1 | Glycosomal | Unknown |
| Tb927.7.310 |  | Glutathione S-transferase, N-terminal domain containing protein, putative | 1 | Dual/multilocalized | Metabolism (Ether lipid synthesis) |
| Tb927.10.14020 | Putative TMEM135 | hypothetical protein, conserved | 10 | Dual/multilocalized | MCS, Metabolism? |
| Tb927.10.8410 | Putative PEX11 family pro- tein | hypothetical protein, conserved | 3 | Glycosomal | Peroxin, organelle proliferation, metabolite transporter |
| Tb927.11.10140 | MDO2 | hypothetical protein, conserved | 0 | Dual/multilocalized | ADP-Ribose binding. Metabolism |
| Tb927.1.5000 |  | hypothetical protein, conserved | 1 (TA) | Glycosomal | Unknown |
| Tb927.11.10020 |  | short chain dehydrogenase/Enoyl- (Acyl carrier protein) reductase, putative | 2 | Dual/multilocalized | Lipid metabolism |
| Tb927.5.1630 | Cytb5-like | Cytochrome b5-like Heme/Steroid binding domain containing protein, putative | 1 | Dual/multilocalized | Metabolism |
| Tb927.8.6830 | Putative KifC3 | Kinesin motor domain/Microtubule binding, putative | 1 | Dual/multilocalized | Organelle transport |
| Tb927.8.6620 | UBA/TS-N | UBA/TS-N domain containing protein, putative | 6 | Dual/multilocalized | Protein quality control, only membrane associated Ubiquitin binding protein |
| Tb927.3.1840 | ECR | 3-oxo-5-alpha-steroid 4-dehydrogenase, putative | 4 | Dual/multilocalized | Lipid metabolism (enoyl-CoA reductase) |
| Tb927.9.2670 | ELMOD2 /POMP3 | ELMO/CED-12 family, putative | 2 | Dual/multilocalized | Unknown |
| Tb927.5.1210 |  | short chain dehydrogenase/KR domain/Enoyl- (Acyl carrier protein) reductase, putative | 2 | Dual/multilocalized | Lipid metabolism, Drug target |
| Tb927.9.11580 | GIM5A |  | 2 |  |  |
| Tb927.9.11640 | Putative PEX11 family protein | hypothetical protein, conserved | 5 | Glycosomal | Peroxisome, organelle proliferation, metabolite transporter |
| Tb927.10.10610 |  | Protein tyrosine phosphatase-like protein, PTPLA, putative | 6 | Dual/multilocalized | Lipid metabolism |
| Tb927.10.9890 |  | hypothetical protein, conserved | 1 | Glycosomal | Unknown |
| Tb927.2.4130 | ECH | Enoyl-CoA hydratase/isomerase/3-hydroxyacyl-CoA dehydrogenase, NAD binding domain/3-hydroxyacyl-CoA dehydrogenase, C-terminal domain containing protein, putative | 1 | Dual/multilocalized | Lipid metabolism |
| Tb927.8.1580 |  | Sulfotransferase domain containing protein, putative | 1 (TA) | Dual/multilocalized | Metabolism/Drug resistance? |
| Tb927.9.10310 | MCP11 | Mitochondrial carrier pro- tein, putative | 1 | Dual/multilocalized | Metabolite transporter |
| Tb927.3.5090 | TXN2 | Redoxin/AhpC/TSA family/Thioredoxin/Thioredoxin-like, putative | 1 (TA) | Dual/multilocalized | Oxidative stress defense |

TA Tail-anchored protein
\*TMD Not predicted by Phobius/TMHMM, 1 TM close to the N-terminus based on orthology.

Further, we performed immunofluorescence microscopy analysis to assess the localization of the GFP-tagged shortlisted proteins to glycosomes of *T. brucei*. Except for *Tb*PMP4, N-terminal GFP tagging was applied since the proteins contained a C-terminal TMD or a putative C-terminal PTS1 like signal. *Trypanosoma* stable cell lines (PCF) encoding various tetracycline-inducible GFP tagged constructs, i.e., PMP4-GFP, GFP-TA1, GFP-TA2, and GFP-MDo2 were induced with 1 µg/mL tetracycline or treated with DMSO alone as negative control. Expression of tetracycline-inducible GFP-tagged constructs was confirmed by immunoblotting. The analysis showed that the tagged proteins migrate at their predicted molecular weight, except GFP-TA2, which runs at ∼70 kDa (and shows a pattern of bands that may represent post-translationally modified forms) **(Fig. S4b)**. Glycosomal localization was assessed by the analysis of colocalization of the fluorescent GFP-fusion of the above constructs with the glycosomal marker enzyme aldolase, which was monitored by immunofluorescence microscopy. All expressed GFP-tagged PMP constructs (PMP4, TA1, TA2, and MDo2) colocalized with the glycosomal marker, implying that the new proteins are indeed glycosomal (**Fig. 3c**). Pearson’s correlation coefficients were used to quantify the colocalization, and the results indicate a significant correlation for GFP-tagged PMPs with aldolase (**Fig. 3d**). Expression of the GFP-tagged PMP4 and the PEX15 candidate 1 and 2, the glycosomes appeared to cluster. Such a clustering has also been observed previously upon expression of other glycosomal membrane proteins, like PEX11 [54, 65] and PEX16 of *T. brucei* [66]. Further, intact *Trypanosoma* cells were sequentially treated with digitonin (to permeabilize plasma membrane and release cytosol) and alkaline carbonate buffer to investigate whether the shortlisted proteins are indeed carbonate resistant integral membrane proteins of glycosomes. Immunoblot analysis shows that all four tested GFP-tagged proteins are carbonate resistant like the glycosomal integral membrane protein markers PEX11 and GIM5 (**Fig. 3c**, right panel), whereas GFP is soluble in the cytosolic fraction along with the cytosolic marker enolase. Taken together, these results show that the inventory’s shortlisted PMP candidates are indeed glycosomal integral membrane proteins.

### Identification of PEX15 in trypanosomatids

A major aim of this study was the identification of the trypanosomal PEX15, a tail-anchored peroxisomal protein in other species, which shows low sequence conservation but high structural similarity [28, 67]. With this as starting information, the two tail-anchored proteins TA1 and TA2 identified in our glycosomal membrane protein inventory were modeled using the AlphaFold2 server and compared with the yeast PEX15 structure. Of the two *Trypanosoma* PEX15 candidates, TA2 lacks structural similarity with *Sc*PEX15 and was not further considered as PEX15 candidate (**Fig. 4a**). However, TA1 shows structural similarity with *Sc*PEX15, with a common N-terminal helical bundle that is known to interact with PEX6 across organisms. In the following, TA1 will be referred to as *Trypanosoma* PEX15 (*Tb*PEX15).

**Fig. 4:**
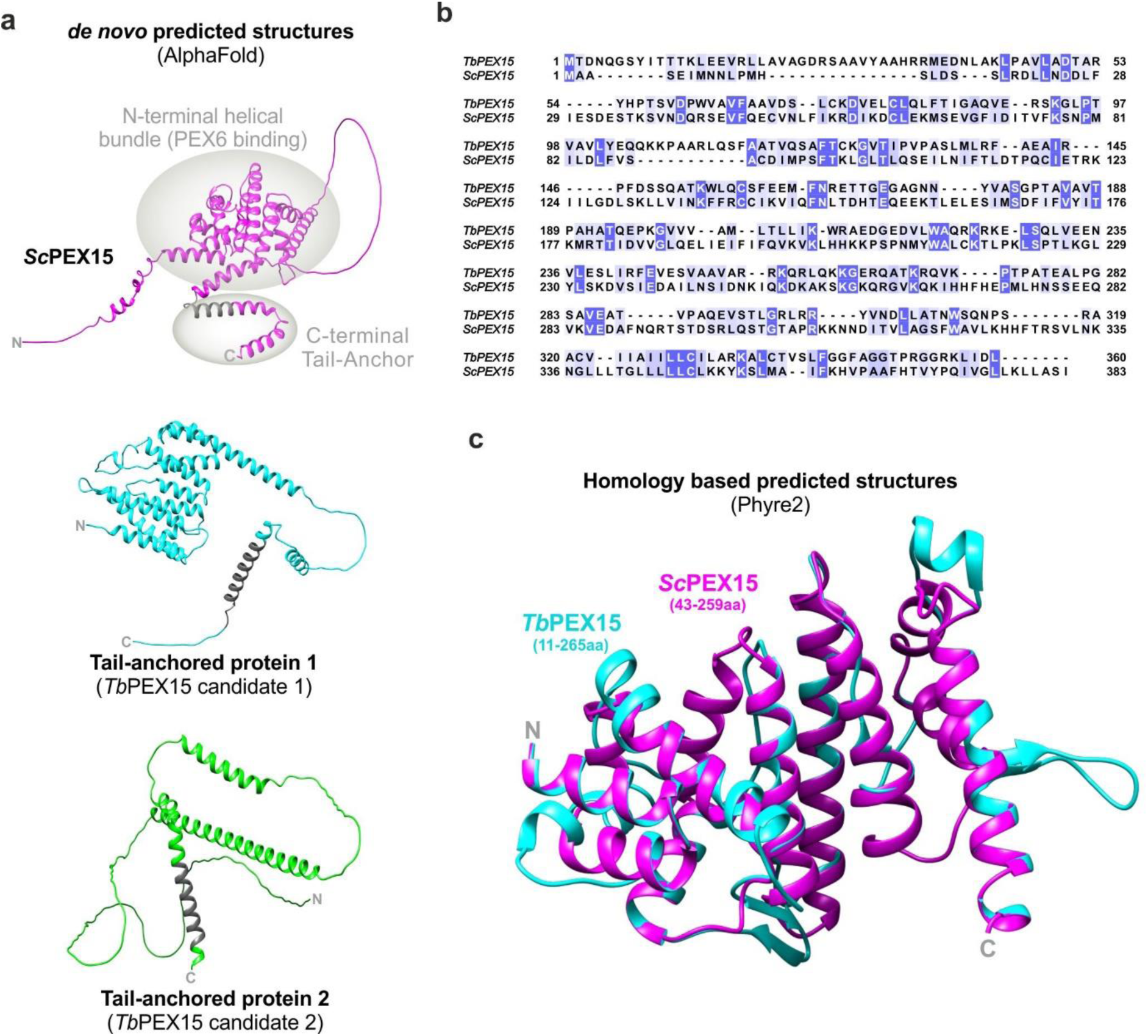
Identification of the trypanosomal PEX15 by comparative modeling. **a** AlphaFold predicted structures of *Saccharomyces cerevisiae* (*Sc*) PEX15 (magenta, left panel), and *T. brucei* (*Tb*) PEX15 candidates TA1 and TA2, of which candidate 1 (middle panel) shows high structural similarity with a common N-terminal helical bundle that for the yeast protein is known to interact with PEX6. A predicted C-terminal TMD that is characteristic for the tail-anchored PEX15 is marked in grey. **b** Multiple sequence alignment of *Trypanosoma* (TriTrypDB ID: <u>Tb927.10.1860</u>) and *Sc*PEX15 proteins. The sequence conservation is colored according to the percentage identity with conservation threshold of 30%. **c** Structural comparison of the putative *Tb*PEX15 modelled using Phyre2 tool (represented in cyan) with the crystal structure of *Sc*PEX15 (PDB Id: 5VXV, represented in magenta) as a template.

As expected, sequence alignment of *Trypanosoma* and yeast PEX15 proteins using the Clustal Omega tool only revealed a low sequence identity of 11.94% and a similarity of 23.05% between the proteins (**Fig. 4b**). A multiple alignment with its counterpart from yeast, human, and plant confirmed the low sequence identity ranging from about 12% to 16% and ∼23-26% sequence similarity (**Figs. S5a, b**). Additional structural comparison of *Trypanosoma* PEX15 by comparative modeling using the Phyre2 tool with the available crystal structure of the soluble part of yeast PEX15 [68] revealed structural similarity (**Fig. 4c**). The modelled structure of PEX15 using the AlphaFold2 server and Phyre2 were validated using the Ramachandran plot, where ≥98 % of the residues were in the allowed region, indicating the quality of the predicted structures (**Fig. S6**). Thus, structure analysis by *de novo* as well as homology modeling strongly suggests that the identified TA1 protein is indeed *Trypanosoma* PEX15.

Furthermore, we compared the predicted 3D structure of *Trypanosoma* PEX15 with its counterparts from yeast, human, and plants (**Figs. S5c, d**). In line with the literature [67], all the structures mentioned above share a feature of N-terminal helical bundle known to interact with PEX6 across different organisms, which provides additional evidence for the identified *Trypanosoma* PEX15. The syntenic orthologs of *Tb*PEX15 are present in all *Trypanosoma* species. However, due to the lack of synteny for *Leishmania* species, we identified the *Leishmania* PEX15 ortholog using a BLAST search with the *Tb*PEX15 sequence. We retrieved the sequences of the *Tb*PEX15 and its counterparts in *T. cruzi* and *L. donovani* from TriTrypDB and aligned them. The *Tb*PEX15 protein shares 33.05% and 18.24% identity with its counterparts in *T. cruzi* and *L. donovani*, respectively, while the similarity is 43.61% and 28.07%, respectively (**Fig. S7a**). We also compared the predicted structure of *Trypanosoma* PEX15 with its counterparts in *T. cruzi* and *L. donovani* obtained from the TriTryp specific AlphaFold database [69] (**Fig. S7b**). The predicted structures of PEX15 in *T. cruzi* and *L. donovani* feature a shared N-terminal helical bundle, as observed for *Sc*PEX15, *Hs*PEX26, and plant APEM9, facilitating the identification of a pan-trypanosomatid PEX15.

### Interaction between PEX15 and PEX6

Pex15 in yeast, PEX26 in human, and APEM9 in plants are membrane anchor proteins for the PEX1-PEX6-complex of the peroxisomal exportomer, which is bound via interaction of the anchor protein PEX6 [67]. Accordingly, the identified *Trypanosoma* PEX15 was tested for its interaction with *Tb*PEX6. To this end, a yeast two-hybrid assay (Y2H) was performed and *Tb*PEX15 constructs, i.e., full-length_1-360aa_ and ΔTMD_1-320aa_ (lacking the transmembrane domain) fused to GAL4-BD were assessed for their interaction with full-length *Tb*PEX6 (**Fig. 5a**). *Tb*PEX15 full-length_1-360aa_ showed a clear interaction with full-length *Tb*PEX6 in the plate-based assay (sensitive), whereas in the liquid assay (quantitative), the interaction appeared weak compared to the positive control. This could be due to the presence of the TMD in *Tb*PEX15, which might impede its translocation into the nucleus and subsequent activation of the GAL promoter. Accordingly, *Tb*PEX15 ΔTMD_1-320aa,_ i.e., the cytoplasmic part exhibited clear and strong interactions with full-length *Tb*PEX6, in both plate-based and liquid assay. The controls with either of the fusion proteins (GAL4-AD/BD) confirmed that there is no auto-activation when the respective fusions with GAL4-AD/BD were tested alone.

**Fig. 5:**
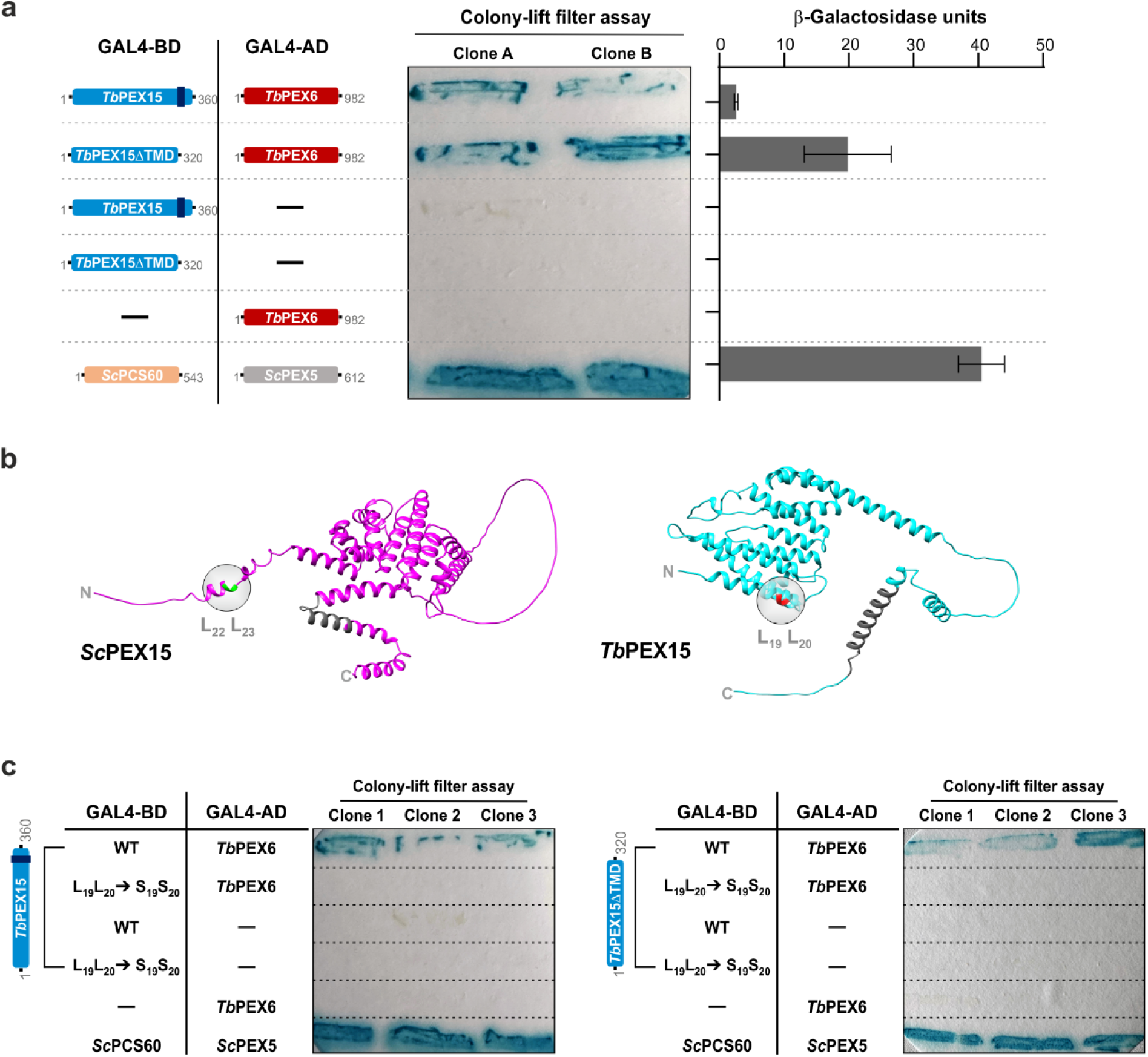
Validation of *Tb*PEX15: Analysis of its interaction with *Tb*PEX6. **a** Y2H interaction analysis of *Tb*PEX15 (full-length or ΔTMD; fused to GAL4 activation domain) and *Tb*PEX6 (fused to GAL4 binding domain) by Colony-lift filter assays (middle panel) and liquid ONPG assays (right panel), with yeast PEX5-PCS60 as a positive control. *Tb*PEX15 (full-length as well as ΔTMD) showed a clear interaction with *Tb*PEX6. The presented β-galactosidase activity units represent the average values of technical replicates from three different biological clones, with error bars indicating the mean and standard deviations. **b** *Tb*PEX15 contains two consecutive leucine residues (labelled in red, right panel) similar to *Sc*PEX15 (labelled in green, left panel) for which the L_22_ to serine mutation is known to abolish the interaction with *Sc*PEX6. The C-terminal TMD is marked in grey. **c** Mutational analysis of the *Tb*PEX15-*Tb*PEX6 interaction in Y2H assay. Mutation of L_19_L_20_ to Serine residues abolishes the interaction of full length (left panel) as well as of the ΔTMD (right panel) variant of *Tb*PEX15 with *Tb*PEX6. The assay was performed in three replicates with three different clones.

To further analyze this putative interaction, we introduced mutations in *Trypanosoma* PEX15 and assessed their effect on the interaction with PEX6. In yeast, exchange of leucine at position 22 to serine of *Sc*PEX15 abrogates the interaction with *Sc*PEX6 [27]. Close inspection of the corresponding region revealed two leucine residues, including L_22_ that are located in the first N-terminal helical region of *Sc*PEX15 (**Fig. 5b**, left panel). Interestingly, there are two consecutive leucine residues at position 19 and 20 (L_19_L_20_) in the first N-terminal helical region of the *Trypanosoma* PEX15 (**Fig. 5b**, right panel). Accordingly, these residues were replaced with serine in *Tb*PEX15 and the effect on the interaction with *Tb*PEX6 was assessed using the Y2H assay. Mutation of leucine residues to serine i.e., L_19_L_20_ to S_19_S_20_ in *Tb*PEX15 (full length and ΔTMD) abolished the interaction with *Tb*PEX6 (**Fig. 5c**). Similar to wildtype *Tb*PEX15, the GAL4-BD fusions of mutated *Tb*PEX15 (full-length and ΔTMD) did not show any autoactivation.

### *Tb*PEX15 is essential for glycosome biogenesis and trypanosomatid parasite survival

To investigate whether *Tb*PEX15 is essential for *T. brucei* parasite survival, we performed a knockdown experiment using RNA interference (RNAi) in the clinically relevant bloodstream form of the parasites. A stable cell line with a genomically integrated tetracycline-inducible RNAi construct targeting the coding region of *Tb*PEX15 was generated. Induction with tetracycline produces double-stranded stem-loop RNA, which leads to degradation of PEX15 mRNA and thus depletion of *Tb*PEX15 expression. Growth of cells treated with tetracycline (Tet, 1 μg/mL) or an equivalent volume of DMSO that served as a negative control was monitored for four days (**Fig. 6a**). Growth defects were observed in the tetracycline-induced RNAi cell line from day 1 onwards, with a ∼40% reduction in cell number compared to DMSO-treated cells. A severe growth defect of the RNAi-induced cultures was evident at day 2, where dead cell debris was clearly seen upon microscopic inspection. To confirm that the growth defect was indeed correlated with *Tb*PEX15 knockdown, we analyzed *Tb*PEX15 protein expression levels (**Fig. 6b**). Wild-type cells treated with DMSO (Tet **-**) or Tet (Tet **+**), and PEX15 RNAi cell line treated with DMSO (Tet **-**) served as negative controls for the experiment, while tubulin served as loading control in the immunoblot analysis. The results show a significant and specific reduction in PEX15 protein levels upon RNAi induction (**Fig. 6b**, Tet +, quantification shown in upper panel of **Fig. 6c**).

**Fig. 6:**
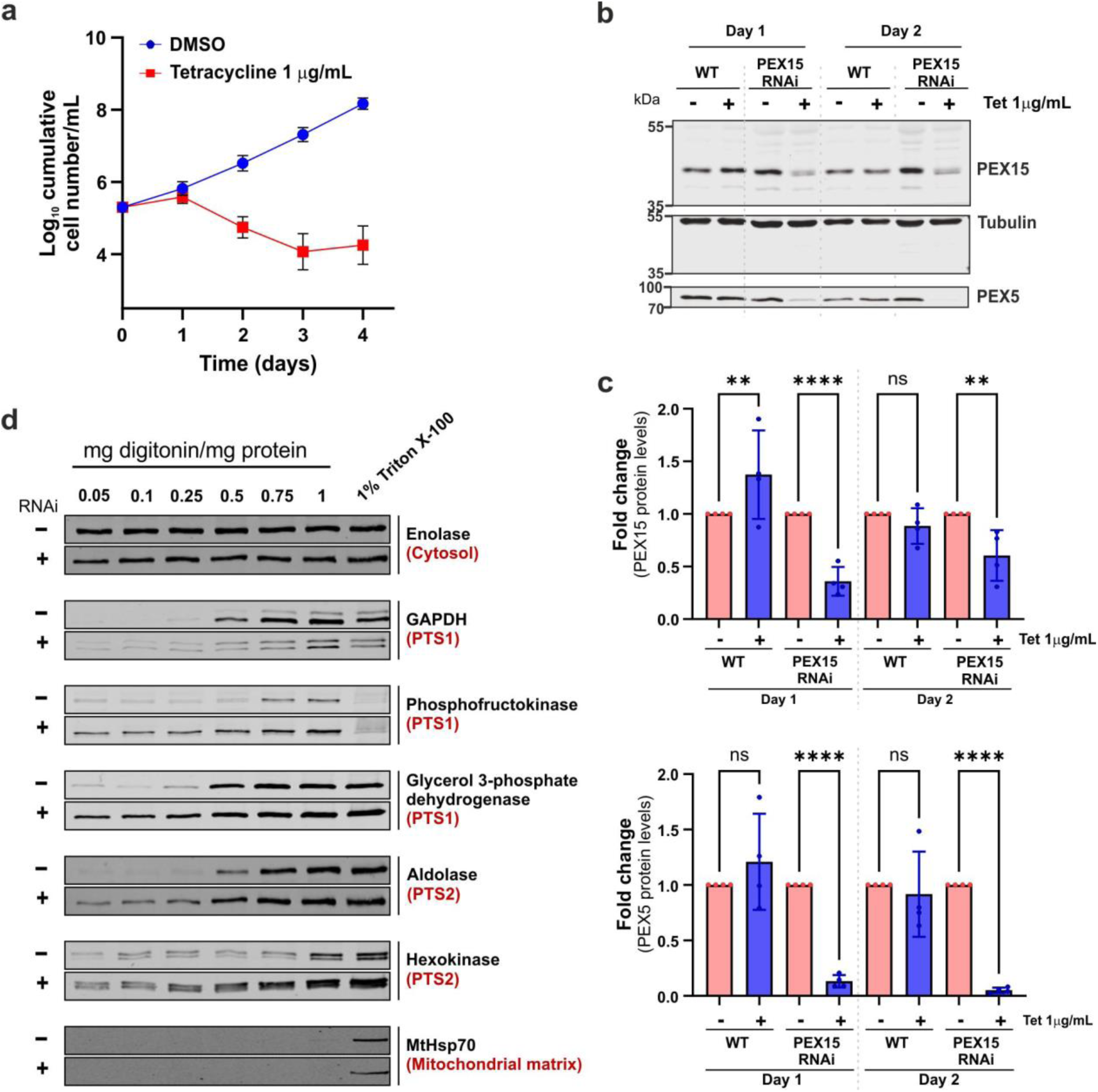
PEX15 is essential for *Trypanosoma* glycosome biogenesis. **a** Cumulative growth curves for uninduced (DMSO) and induced (Tetracycline) bloodstream form PEX15 RNAi cell lines demonstrate that *Tb*PEX15 is essential for parasite survival (upper panel). The experiment was conducted in biological triplicates, and error bars represent the standard error of the mean. **b** Depletion of endogenous PEX15 and PEX5 upon PEX15 RNAi: Immunoblot analysis of total cell lysates from wildtype and PEX15 RNAi cells, treated with DMSO or tetracycline, was performed using antibodies against PEX15, alpha tubulin (loading control), and PEX5. Steady-state concentrations of PEX5 were significantly decreased upon induction of PEX15 RNAi on day 1 and day 2. **c** the bar graph depicts the fold changes of *Tb*PEX15 and *Tb*PEX5 in wild-type (WT) and *Tb*PEX15 RNAi-inducible cells following tetracycline treatment, compared to their respective controls. α-Tubulin was used as a loading control. Statistical analysis was performed using the one-way ANOVA test across four biological replicates. ****p<0.0001; ***p<0.001; **p<0.01; *p<0.05; ns, not significant. The error bars represent the mean ± standard deviation. **d** RNAi-mediated depletion of PEX15 results in the mislocalization of glycosomal matrix proteins as shown by biochemical fractionation of digitonin-permeabilized cells. BSF parasites treated for 2 days with DMSO (-) or tetracycline (+) for PEX15 RNAi-induction were incubated with increasing digitonin concentrations. Cell suspensions were subjected to centrifugation to separate the soluble fractions, which were further analyzed by immunoblotting using various antibodies as indicated. Cells treated with 1% Triton-X 100 represent complete release of proteins by dissolving all membranes. Enolase served as loading control.

Further, we assessed the impact of *Tb*PEX15 RNAi on PTS1-receptor (PEX5) recycling, since this process requires anchoring of the PEX1-PEX6 complex to the peroxisomal/glycosomal membrane via PEX15. Thus, PEX15 depletion should result in the degradation of PEX5, which was confirmed by significantly reduced PEX5 levels through quantitative immunoblot analysis (**Fig. 6b, lower panel;** quantification shown in **Fig. 6d, lower panel**). This result is consistent with PEX5 instability in PEX1-depleted cells of *T. brucei* [32] and supports the identity and function of the so far unknown *Tb*PEX15 presented in this work.

A common feature of the depletion of a peroxin in *T. brucei* is the disruption of glycosome biogenesis, leading to the mislocalization of glycosomal matrix enzymes to the cytosol, which in turn causes parasite death [13, 70]. We investigated the mislocalization of glycosomal matrix proteins in BSF parasites upon *Tb*PEX15 RNAi. As a clear growth defect was observed at day 2 upon *Tb*PEX15 RNAi-induction, these cells were used for the subcellular localization of the glycosomal PTS1-protein GAPDH and the PTS2-protein aldolase by immunofluorescence microscopy. DMSO-treated cells served as control and show a bright punctate pattern for the glycosomal markers, indicative for a functional import of the marker proteins into glycosomes (**Fig. S8**, top panel). The tetracycline-induced RNAi cells exhibited a similar labelling of both GAPDH and aldolase although the signals were less bright (**Fig. S8**, lower panels). However, mislocalization of the glycosomal enzymes to the cytosol was not discernible. Since the RNAi-induced parasites still contain previously imported glycosomal enzymes, this may mask a diffuse cytosolic staining in microscopy images and hamper a quantification of the import defect by this technique. Therefore, we performed a biochemical fractionation assay, which provides a quantitative method for assessing the mislocalization of glycosomal matrix proteins to the cytosol. To this end, PEX15 RNAi cells were treated for 2 days with DMSO or tetracycline (RNAi-induced). Further, these cells were harvested and treated with increasing digitonin concentration. Following centrifugation, the supernatants were analyzed by immunoblotting using antibodies against markers proteins of the cytosol, glycosomal matrix, glycosomal membrane, and mitochondrial matrix. Cells treated with 0.05 mg/mg of protein completely released the cytosolic protein enolase, serving as a loading control (**Fig. 6c**). Higher concentrations of digitonin (0.5 – 1 mg/mg protein) were required to release glycosomal proteins into the supernatant in the DMSO-treated cells. Only a modest amount of glycosomal proteins, such as phosphofructokinase and hexokinase, were released with lower concentrations of digitonin, as described before [66]. However, in PEX15 RNAi-induced cells, both PTS1- and PTS2- containing glycosomal matrix enzymes were well detected in the supernatants already at lowest digitonin concentrations, which also released cytosolic enolase, indicating the mislocalization of glycosomal matrix proteins to the cytosol upon PEX15 RNAi knockdown (**Fig. 6c**, panels labeled as **+**).

Taken together, our data demonstrate that PEX15 function is essential for the biogenesis of glycosomes in the bloodstream form of *T. brucei* cells and thus PEX15 depletion is lethal for *Trypanosoma* parasites.

## Discussion

Peroxisomes are versatile organelles that can adapt their proteome according to cellular needs. Likewise, glycosomes provide metabolic plasticity to the trypanosomatid parasites, which infect the mammalian host and encounter different metabolic niches. In this study, we established a high confidence inventory of glycosomal membrane proteins of *Trypanosoma* parasites. In-depth quantitative proteomics analysis of highly purified glycosomal membranes and protein correlation profiling across density gradient fractions followed by statistical analysis and classification resulted in the identification of 28 novel glycosomal membrane proteins of high confidence. This dual subcellular proteomics approach is effective in identifying also multi-localized glycosomal membrane proteins while separating them from co-purified components of non-glycosomal localization. Correlation analysis using manually curated list of various *Trypanosoma* organelle markers revealed that 17 of the 28 novel proteins are dually localized, mostly to mitochondria or the ER. Two known mitochondrial outer membrane proteins observed in our inventory are Msp1 and 3-oxo-5-alpha-steroid 4-dehydrogenase [46, 60]. Msp1 (ATAD1 in humans) is an AAA-ATPase, which recently has been shown to localize to both glycosomes and the mitochondrial outer membrane where it plays a role in protein quality control [61].

A structure-based search revealed potential homologs of TMEM135 and TbSDR, a putative acyl/alkyl DHAP reductase. Notably, profiling of *Tb*TMEM135 and *Tb*SDR indicated their dual localization to glycosomes as well as to mitochondria/ER, which is not distinguished in our study. In this aspect, the trypanosomal TMEM135 seems to behave like its human counterpart, which is a multifunctional protein that localizes to both mitochondria and peroxisomes, serving as a critical mediator in the peroxisomal regulation of mitochondrial fission [71–73]. Interestingly, TMEM135 has also been reported to be involved in lipid droplet formation/tethering, fatty acid metabolism, and peroxisomal function [71, 73]. TbSDR is a homologue of ADHAPR, also known as PexRAP (Peroxisomal Reductase Activating PPARγ), a peroxisomal lipid synthesis enzyme that plays a role in adipogenesis. This enzyme is localized to both peroxisomes and the endoplasmic reticulum [74, 75]. Interestingly, in *Trypanosoma brucei*, alkyl/acyl-DHAP reductase activity was found to be associated with glycosomes [55, 76]. However, the gene encoding the alkyl/acyl-DHAP reductase has not yet been identified [77]. Whether this glycosomal enzyme activity can be attributed to TbSDR requires further investigation.

Members of the PEX11-family are abundant integral peroxisomal membrane proteins involved in peroxisome function and biogenesis in yeasts, plants, and mammals [78]. In trypanosomes, three members of this family, *Tb*PEX11 and GIM5a/b, have been identified and it has been shown that they are essential for parasite survival [54, 79]. Here, we identified two new PEX11 family proteins in *T. brucei* by a structure-based approach. For yeast and humans, three PEX11 members have been identified, plants have five members [80]. With now five members of this family, the trypanosomatids in this aspect are similar to plants and further investigations are required to determine the role of these proteins in glycosome function.

Analysis of the *Trypanosoma* PMP4 revealed that it is a homolog of the human peroxisomal membrane protein PXMP4 (also known as PMP24 or PMP4), an evolutionarily highly conserved protein among eukaryotes whose function remains unknown [81, 82]. The *Tb*PMP4 was also reported in the PMP inventory of *T. brucei* [46].

MDo2 is the second macrodomain-like protein of trypanosomatids that has been identified. Previously, macrodomain-like protein, i.e., MDo1, was identified and characterized as ADP-ribose hydrolase in *T. brucei* and *T. cruzi*, which are homologous to human O-acetyl-ADP-ribose deacetylases MacroD1 and MacroD2 [63]. However, the study did not emphasize its subcellular localization. Interestingly, MDo1 contains a PTS1-like signal (-SSL) at its C-terminus and is also reported as glycosomal protein of high confidence [47]. Remarkably, MDo2, which contains a predicted TMD in its N-terminal region and seems to be a bona fide glycosomal membrane protein, also contains a PTS1 (-SKL) at its extreme C-Terminus. So far, such a constellation has only been reported for PEX13.1, an essential protein of the glycosomal protein import machinery [64]. The significance of the PTS1 signal for the topogenesis or function of MDo2 is an exciting question for further investigation. Fluorescence microscopy investigation of the subcellular localization of the N-terminally tagged protein revealed that it is targeted to glycosomes. However, the profiling showed that the protein is dually localized also to mitochondria/ER. It is feasible that the N-terminal tagging might interfere with its targeting. Along this line, MDo2 has been identified in the mitochondrial outer membrane proteome [83]. Accordingly, MDo2 protein is a dually localized protein to glycosomes and mitochondria.

PEX15 of *T. brucei* was identified as a one of the class 1 proteins. Most of the peroxins or PEX proteins in trypanosomes were discovered by their sequence similarity with the known PEX proteins of other organisms. However, identification of *Trypanosoma* PEX15 (i.e. essential anchor of the PEX1-PEX6 complex) by BLAST search failed. Yeast PEX15 and its counterparts from other organisms, like human PEX26 and plant APEM9 share very low amino acid sequence identity, but they all are Tail-anchored proteins and they share a high structural similarity of the cytosol facing domain [28, 67]. As the low sequence similarity could make PEX15 an ideal drug target, its identification was a major aim of this study. In this respect, the two hypothetical tail-anchored proteins present in the Class 1 inventory were identified as promising PEX15 candidates. Our data provide a convincing chain of evidence that one of the newly identified tail-anchored proteins indeed represents PEX15. For PEX15, it could be predicted that 1) it localizes to glycosomes as a tail-anchored membrane protein, 2) its depletion as a peroxin should result in defect in glycosomal protein import and killing of the parasites, and 3) it would interact with PEX6 as membrane anchor for the AAA-complex of PEX1/PEX6. In this report, we confirmed all these assumptions for one of the newly identified proteins, leaving no doubts that it represents the bona fide trypanosomal PEX15. This conclusion is further supported by two additional observations. First, despite a low sequence similarity, the newly identified *Tb*PEX15 shares a striking structural similarity with *Sc*PEX15, *Hs*PEX26, and APEM9 including a common N-terminal helical bundle known to interact with PEX6 (reviewed in [67]). The structural similarity is especially highlighted by homology-based modeling superimposing *Tb*PEX15 with the crystal structure of *Sc*PEX15 [68]. Finally, for components that are required for recycling, it is known that their depletion results in an instability of the PTS1-receptor PEX5. This is because in the absence of recycling PEX5 accumulates at the peroxisomal membrane, gets polyubiquitinated and is then degraded by a quality control pathway named RADAR (Receptor Accumulation and Degradation in the Absence of Recycling) [84–86]. This results in a decreased steady-state level of PEX5, which has also been described for *T. brucei* upon depletion of *Tb*PEX1, a protein involved in receptor recycling [32]. This decreased steady-state level is also observed upon depletion of PEX15, fostering its role as component of the glycosomal import receptor recycling machinery. Taken together, our data are clear in that the identified protein is bona fide PEX15 of *Trypanosoma* parasites. With respect to drug development the low degree of sequence conservation compared to its human counterpart, makes *Tb*PEX15 an attractive target. This finding holds significant therapeutic implications for developing drugs combating *Trypanosoma* parasites.

## Conclusion

Taken together, this study provides a high-confidence protein inventory of glycosomal membranes in *T. brucei*. It includes several unique and parasite-specific uncharacterized proteins, which could be hitherto unknown metabolite transporters or proteins involved in glycosomal biogenesis, membrane contact sites, interorganelle communication, protein quality control, and more. Therefore, this study establishes a strong foundation for future research on the components and unique mechanisms underlying glycosome biogenesis and its functions. This research is essential for developing treatments to combat trypanosomatid diseases. Most importantly, this study identified *Trypanosoma* PEX15, the so far unknown membrane anchor for the glycosomal AAA-complex that is responsible for receptor recycling, which is an essential step in matrix protein import. Considering its low degree of sequence conservation compared to its human counterpart, the *Tb*PEX15 is an attractive target for drug development.

## Methods

### *Trypanosoma* cell culture and transfection

In this study, *Trypanosoma brucei brucei* procyclic form (PCF) 29-13 and bloodstream form (BSF) 90-13 cell lines, co-expressing T7 RNAP and TetR, were used. The BSF cells were cultured in HMI-11 medium [87] at 37 °C in a humidified incubator with 5% CO_2_, while PCF cells were grown in SDM-79 medium [88] at 28 °C. Both cultures were supplemented with 10% fetal bovine serum (FBS). PCF cultures were maintained at 1×10^6^– 30×10^6^ cells/ml, while the BSF cultures were kept in the logarithmic growth phase (below 2 × 10^6^ cells/ml). Transfections were performed with NotI-linearized plasmid constructs, which were stably integrated into the rRNA locus within the genome of the respective cell lines. Clones were selected using the antibiotics as previously described in [66]. Transfection of PCF and BSF cells was performed as described previously in [31] and [66], respectively. The antibiotic concentrations used for the selection of clones with respective plasmid constructs are as follows: PCF cells, pGN1/pGC1 (10 μg/ml Blasticidin); and BSF cells, pGN1/pGC1 (5 μg/ml Blasticidin) or p2T7-177 (2.5 μg/ml Phleomycin). Additionally, G418 (15 μg/ml for PCF cells and 2.5 μg/ml for BSF cells) and Hygromycin (50 μg/ml for PCF cells and 5 μg/ml for BSF cells) antibiotics are used for cell maintenance, in addition to the antibiotics used for selection.

### Cell lysis, subcellular fractionation, and glycosome isolation

Procyclic *Trypanosoma* cells (approx. 1.9 × 10^10^ cells per experiment) were harvested by centrifugation for 10 min at 1,500 g (Rotor SX4400, Beckman Coulter) and 4 °C and were resuspended in homogenization medium (250 mM sucrose, 25 mM Tris-HCl, 25 mM NaCl, 1 mM EDTA, pH 7.6, supplemented with 1x protease inhibitor cocktail (PIC; Roche/Merck Cat. No. 4693132001). Cells were ruptured with silicon carbide in a pre-cooled mortar, followed by differential centrifugation of the homogenate, which yields an organelle-enriched fraction [52]. This organelle-enriched fraction generally contains mitochondrial-derived vesicles, lysosomes, glycosomes, and endoplasmic reticulum (ER) vesicles. Subsequent density gradient centrifugation (20 – 46% OptiPrep^TM^-sucrose solution) of the organelles was performed to purify intact glycosomes, as outlined in the previous work [52], which results in the separation of different cellular components according to their buoyant density. Following centrifugation, a clear separation of the organelles into two bands was visible, with the lower one containing intact glycosomes [44, 47], while the other upper band contains organelles such as mitochondria, ER, and others [44, 60] (see **Fig. 1a**). Fractions of 0.5 mL each were collected by puncturing the tube at the bottom. Subsequently, aliquots of each fraction were analyzed by SDS-PAGE and immunoblotting. The glycosomal peak fractions 12 and 13 (**Fig. 1b**) were pooled, and glycosomes were concentrated by centrifugation at 20,800 g and 4 °C for 30 min (Rotor F45-30-11, Eppendorf). The resulting pellet was then used for the enrichment of glycosomal membrane proteins.

### Enrichment of glycosomal membrane proteins

Glycosomal membrane proteins were enriched as described before [52]. In brief, the glycosomal pellet was resuspended in 350 μL of low-salt buffer (5 mM Tris-HCl, pH 7.6, 1 mM EDTA, 1X PIC) and incubated for 30 min at 4 °C. Centrifugation for 30 min at 96,000 g and 4 °C (Rotor TLA-100, Beckman Coulter) results in a supernatant, containing soluble proteins including the glycosomal matrix proteins, and a pellet, containing membrane-associated proteins. The pellet was resuspended in 300 μL of alkaline carbonate buffer (100 mM Na_2_CO_3_, pH 11.5) and incubated for 60 min at 4 °C. The suspension was then layered onto a 120-μL sucrose cushion (250 mM sucrose, 100 mM Na_2_CO_3_, pH 11.5) and centrifuged for 60 min at 150,000 g and 4 °C (Rotor TLA-100, Beckman Coulter). The pellet was again subjected to alkaline carbonate extraction to minimize the contamination of matrix proteins. The supernatant fractions of this procedure contain soluble matrix proteins and peripheral membrane proteins, while the carbonate-resistant pellet represents the sedimentation of integral membrane proteins. The resulting sediment was resuspended in a carbonate buffer containing sucrose (250 mM sucrose, 100 mM Na_2_CO_3_, pH 11.5). After each centrifugation step, aliquots of supernatant and pellet were taken for immunoblot and MS analysis.

### Enrichment of membrane proteins from density gradient fractions

Organelles present in fractions #11 to #22, collected from a 20 – 46% OptiPrep^TM^ sucrose gradient as described above, were collected by centrifugation (20,800 g, 30 min, 4 °C) and resuspended in 160 μL of alkaline carbonate buffer (100 mM Na_2_CO_3_, pH 11.5). Following incubation for 60 min at 4 °C, samples were layered onto a 60-μL sucrose cushion (250 mM sucrose, 100 mM Na_2_CO_3_, pH 11.5) and were centrifuged for 60 min at 150,000 g and 4 °C (Rotor TLA-100, Beckman Coulter). The pellets were resuspended in carbonate buffer containing sucrose (250 mM sucrose, 100 mM Na_2_CO_3_, pH 11.5) and analyzed by immunoblotting and MS (**Figs. 1e, f**).

### LC-MS sample preparation

Proteins (20 µg per sample) were acetone-precipitated and subsequently resuspended in urea buffer consisting of 8 M urea and 50 mM ammonium bicarbonate (ABC). Cysteine residues were reduced and alkylated by sequential incubation with 5 mM Tris (2-carboxyethyl) phosphine dissolved in 10 mM ABC (30 min at 37 °C) and 55 mM iodoacetamide/10 mM ABC (30 min at room temperature in the dark). Reactions were terminated by adding dithiothreitol (25 mM final concentration). Samples were first diluted to 4 M urea using 50 mM ABC to digest proteins with LysC (4 h at 37 °C) and subsequently to 1 M urea for digestion with trypsin (overnight at 37 °C). Proteolytic reactions were stopped by adding trifluoroacetic acid (TFA) to a final concentration of 1% (v/v). Peptides were desalted using StageTips [89] and dried under vacuum.

### LC-MS analysis

Peptide mixtures, reconstituted in 0.1% TFA, were analyzed by nano HPLC-ESI-MS/MS using an UltiMate 3000 RSLCnano HPLC system (Thermo Fisher Scientific, Dreieich, Germany) online coupled to an Orbitrap Elite hybrid mass spectrometer (Thermo Fisher Scientific, Bremen, Germany). The RSLC system was equipped with PepMap C18 precolumns (5 mm x 300 µm inner diameter, Thermo Scientific) and an analytical C18 reversed-phase nano LC column (nanoEase M/Z HSS C18 T3; 250 mm length, 75 mm inner diameter, 1.8 mm particle size, 100 Å packing density; Waters A binary solvent system consisting of 4% (v/v) dimethyl sulfoxide (DMSO)/0.1% (v/v) formic acid (FA) as solvent A and 30% (v/v) acetonitrile (ACN)/48% (v/v) methanol/4% (v/v) DMSO/0.1% (v/v) FA as solvent B was employed for peptide separation. Peptide mixtures equivalent to 1 µg of protein were loaded, washed and preconcentrated on the pre-column for 5 min using solvent A and a flow rate of 10 µL/min. A gradient was then applied at a flow rate of 300 nL/min ranging from 1–7% solvent B in 5 minutes, 7–50% B in 245 min, 50-95% B in 85 min, and 5 min at 95% B. Eluted peptides were transferred to a fused silica emitter for electrospray ionization facilitated by a Nanospray Flex ion source with DirectJunction adaptor (Thermo Fisher Scientific), applying a spray voltage of 1.8 kV and a capillary temperature of 200 °C. MS/MS data were acquired in data-dependent mode using the following parameters: MS precursor scans at *m/z* 370–1700 with a resolution of 120,000 (at *m/z* 400); automatic gain control (AGC) of 1 × 10^6^ ions; a maximum injection time (IT) of 200 ms; a TOP20 method for low-energy collision-induced dissociation of multiply charged precursor ions with a normalized collision energy of 35%, an activation q of 0.25, and an activation time of 10 ms. The AGC for MS/MS scans was set to 5 × 10^3^ with a maximum IT of 150 ms and a dynamic exclusion time of 45 s.

### MS data analysis

For the analysis of mass spectrometric data, the MaxQuant software package (version 2.0.2.0) [90] with its integrated Andromeda search engine [91] was employed. MS/MS data were searched against all protein sequences for *T. brucei* Lister 427 downloaded from the TriTrypDB (release 48; https://tritrypdb.org/tritrypdb/app). Protein identification was performed using MaxQuant default settings (including carbamidomethylation of cysteine residues as fixed modification and N-terminal acetylation and methionine oxidation as variable modifications), with the following exceptions: at least one unique peptide was required for protein identification, ‘Trypsin/P’ and ‘LysC/P’ were set as proteolytic enzymes, and a maximum of three missed cleavages was allowed. The options ‘match between runs’ and ‘iBAQ’ (i.e., intensity-based absolute quantitation) (Schwannhäusser *et al.*, 2011; Tyanova *et al.*, 2016) were activated.

iBAQ intensities were used to establish protein abundance profiles across the fractions of the glycosomal membrane protein enrichment experiment (i.e., total, SN0, SN1, SN2 and MP; n = 3) or carbonate-resistant membrane fractions of the density gradient fractions (fractions F11 - F22; n = 1) for proteins that were identified in at least two out of the five fractions of glycosomal membrane protein enrichment experiments (per replicate) or two out of the twelve membrane fractions of the density gradient, respectively. For data processing, analysis as well as visualization, the autoprot Python module [92] was used. Briefly, for each replicate, iBAQ values for individual proteins were log-transformed, missing values were imputed row-wise by randomly drawing values from a downshifted distribution across rows and normalized to the highest value measured for the protein across all fractions, which was set to one. Information about proteins identified in glycosomal membrane protein enrichment experiments and carbonate-resistant pellets of density gradient fractions are provided in **File S1**.

We used protein correlation profiling to define the glycosomal membrane proteome including new glycosomal membrane proteins. Abundance profiles of proteins detected in glycosomal membrane protein enrichment experiments were correlated replicate-wise with the mean profile of selected glycosomal membrane marker proteins (see **File S1** for information about marker proteins). To this end, the Pearson’s correlation was calculated using the ‘pearsonr’ function from the ‘scipy.stats’ package in Python. The rank sum method (Breitling & Herzyk, 2005; Buitinck et al., 2013; Pedregosa et al., 2011), implemented in the R package ‘RankProd’ (version 3.11) (Del Carratore et al., 2017), was then used to calculate the mean correlation and a p-value for each protein. Proteins with a mean correlation of > 80% to the mean profile of the glycosomal membrane marker proteins were considered as putative glycosomal membrane proteins and divided into two classes based on the p-value (p-value of < 0.01 for class 1, p-value ≥ 0.01 for class 2). Furthermore, the abundance profiles of all candidate proteins across the fractions of the density gradient were compared manually with the mean profiles of glycosomal and mitochondrial/ER marker proteins to remove coenriched contaminants and to enable the detection of glycosomal membrane proteins with additional subcellular localization. Profiles for all proteins present in the preliminary list of putative glycosomal membrane proteins are shown in **File S2**. A Jupyter notebook providing documentation of the analysis pipeline and statistical tools used is available at https://github.com/ag-warscheid/Tb_GMP_Inventory.

#### Cloning

Yeast and *Trypanosoma* expression plasmid constructs and cloning strategies are listed in **Table S1**, and oligonucleotide sequences are listed in **Table S2**. GFP-*Tb*PEX15 was cloned by the FastCloning (In-fusion cloning) method, as described in [93]. Mutations in *Tb*PEX15_1-360aa_ and *Tb*PEX15 ΔTMD_1-320aa_ were generated by overlap extension PCR. Sequences of the constructs, mutations, and gene fragment deletions were verified for all constructs by automated Sanger sequencing.

### Microscopy

*Trypanosoma* stable cell lines (PCF) encoding various tetracycline-inducible GFP tagged constructs [GFP, Peroxisomal membrane protein 4 (PMP4)-GFP, GFP-TA1, GFP-TA2, and GFP-Macro domain 2 (MDo2)] were induced with 1 µg/mL tetracycline or treated with DMSO alone as negative control. Cells were sedimented and fixed by resuspension in 4% paraformaldehyde in PBS (phosphate-buffered saline, pH 7.4) at 4 °C for 15 min. Fixed *Trypanosoma* cells were processed for imaging as described previously [65]. The cells were stained with α-aldolase antibody (1:500) and α-GAPDH (1:500), which are glycosomal markers. Glycosome stained cells were visualized and imaged (five Z-stacks) with a Carl Zeiss microscope using Zen 3.6 software (blue edition). All the images were processed by the deconvolution method and were merged. Further, the processed images were analyzed using Zeiss Zen 3.2 software (blue edition).

### Cell fractionation

For cell fractionation, *Trypanosoma* cells (PCF) encoding GFP tagged constructs (GFP, GFP-PTS1 [18], PMP4-GFP, GFP-TA1, GFP-TA2, and GFP-MDo2) were induced for expression with tetracycline (1 µg/mL). Cells expressing the protein of interest were treated with digitonin at a final concentration of 0.1 mg/mg of protein for 3-5 min at 37 °C. Subsequently, the samples were centrifuged at 20,800 g for 15 min at 4 °C (Rotor F45-30-11, Eppendorf), yielding a cytosol enriched fraction and an organellar pellet. The organellar pellet was further treated with alkaline carbonate buffer (100 mM Na_2_CO_3_, pH 11.5) for 60 min at 4 °C. In the following, the suspension was layered on a cushion (250 mM sucrose, 100 mM Na_2_CO_3_, pH 11.5) and centrifuged for 60 min at 4 °C (150,000 g, Rotor TLA-100, Beckman Coulter). The resulting supernatant contains released matrix and peripheral membrane proteins, while the carbonate-resistant pellet contains integral membrane proteins. Sample collected from each step were analyzed by immunoblotting.

### Bioinformatics analysis and structure modelling

The protein sequences of Human PEX26, Yeast PEX15, and Plant APEM9, which are all orthologs of PEX15, were obtained from the UniProt database with IDs Q7Z412 (*H. sapiens*), Q08215 (*S. cerevisiae*), and Q8W4B2 (*Arabidopsis thaliana*). Additionally, the Human PXMP4 protein sequence was also obtained from the UniProt database (ID: Q9Y6I8). The parasite protein sequences of identified PEX15 (Tail-anchored protein 1) were obtained from the TriTrypDB database with accession IDs Tb427_100021800 (*T. brucei*), TcCLB.508059.20 (*T. cruzi*), and LdBPK_210110.1 (*L. donovani*). The TriTrypDB IDs of other *T. brucei* glycosomal membrane proteins characterized here are PMP4 (Peroxisomal membrane protein 4-Tb427_090008200), Tail-anchored protein 2 (Hypothetical protein-Tb427_010033000) and the MDo2 (Macrodomain 2-Tb427_110110000). Transmembrane segments were predicted using the Phobius web tool (https://www.ebi.ac.uk/Tools/pfa/phobius/) [94]. The domain architecture of the proteins was predicted with the InterPro Scan (https://www.ebi.ac.uk/interpro/) [95]. Multiple sequence alignment was performed using the Clustal Omega tool (https://www.ebi.ac.uk/Tools/msa/clustalo/) [94] and visualized using Jalview (v 2.11.0) software with a percentage identity color scheme. Percentage identity and similarity matrix of proteins were calculated using the SIAS tool with BLOSUM62 matrix (http://imed.med.ucm.es/Tools/sias.html).

The predicted 3D structure of proteins was obtained from the AlphaFold protein structure database [96, 97]. Since the structures of proteins from the *T. brucei* 427 strain were not available in the AlphaFold database, these proteins were modeled using the AlphaFold2 server [96, 98]. The protein IDs modelled using the AlphaFold2 server are Tb427_100021800 and Tb427_010033000. Additionally, *Tb*PEX15 (Tb427_100021800) was also modeled using the Phyre2 tool (One 2 One threading) with the crystal structure of *Sc*PEX15 (PDB Id: 5VXV) as a template (http://www.sbg.bio.ic.ac.uk/phyre2) [68, 99]. The structures modeled using the AlphaFold2 server and Phyre2 tool were validated with the Ramachandran plots generated using the PROCHECK tool from the Saves server (https://saves.mbi.ucla.edu/) [100]. Additionally, 3D structures of *Trypanosoma* proteins (TREU927) were obtained from the TriTryp AlphaFold database (http://wheelerlab.net/alphafold/) [69]. The structures obtained from the TriTryp AlphaFold server include putative *Tb*TMEM135 (Tb927.10.14020), putative short-chain dehydrogenase (Tb927.11.10020), *Tb*PEX11 (Tb927.11.11520) and putative *Tb*PEX11 family protein (Tb927.10.8410 and Tb927.9.11640). To identify homologous proteins, the Foldseek server (https://search.foldseek.com/search) was employed to query the predicted structures, applying a taxonomic filter for *Homo sapiens* [62]. All structures were analyzed and visualized using UCSF Chimera software 1.17.3 (https://www.rbvi.ucsf.edu/chimera) [101].

### Yeast two-hybrid analysis (Y2H)

*S. cerevisiae* wild-type strain PCY2 was grown in double dropout SD synthetic media (without tryptophan and leucine). Yeast cells were transformed by the traditional Lithium-acetate method [102]. Y2H studies were based on the Yeast protocols handbook (Clontech, Protocol No. PT3024-1, Version No. PR742227). Full-length or truncation (ΔTMD) of *Tb*PEX15 with and without mutations were cloned in the pPC97 vector containing GAL4-DNA Binding Domain (BD) as described in **Table S1**. Co-transformation of various two-hybrid plasmids, i.e., BD-*Tb*PEX15 and AD-*Tb*PEX6 [32] constructs, were performed in the Wildtype PCY2 strain. The clones were selected on SD synthetic medium without tryptophan and leucine. A filter-based β-galactosidase assay and liquid culture assay using ONPG were performed in three replicates as described in the Yeast protocols handbook (Clontech). The interaction between GAL4-AD and GAL4-BD fused to *Sc*PEX5 [103] and *Sc*PCS60 [104], respectively, served as a positive control for the study. The GAL4-AD fusion of *Tb*PEX6 and the various GAL4-BD fusions of *Tb*PEX15 (full-length and ΔTMD). WT/mutants were also tested for autoactivation.

### RNA interference and digitonin fractionation

The double-stranded (stem-loop) *Tb*PEX15 RNAi construct was cloned into a *Trypanosoma* expression plasmid p2T7-177 (tetracycline-inducible system), as described in section ‘cloning’. It was then genomically integrated into the BSF *Trypanosoma* cells by transfection, as described in section ‘Trypanosoma cell culture and transfection’. With positive clones, a survival analysis was performed. RNAi was induced upon tetracycline (1 μg/mL) addition to the transfected *Trypanosoma* cells, cultured at 2 × 10^5^ cells/mL density. DMSO treated cells served as a negative control for the study. Following the incubation for 24 h, the cell count was recorded, and the cells were diluted back to the 2 × 10^5^ cells/mL with the addition of tetracycline. The culture with a density below 2 × 10^5^ cells/mL was incubated further without dilution and addition of tetracycline [34]. Growth of the transfected cell line treated with DMSO, or tetracycline was monitored for four days. The experiment was performed in three biological replicates. The cumulative cell number was calculated from cell counts recorded every 24 h and the cumulative growth curve on a logarithmic scale was plotted using GraphPad Prism 10. The DMSO and RNAi-induced cells were harvested on day 1 and day 2, and the cell lysates corresponding to 2 million cells per well were analyzed by immunoblotting with anti-*Tb*PEX15 (for assessing RNAi specificity and efficiency) and anti-*Tb*PEX5 antibodies. In addition, the day 2 samples were also processed for immunofluorescence microscopy as described in section ‘Microscopy’ and biochemical fractionation of cells using digitonin to assess the effect on glycosomal protein import.

Biochemical fractionation using digitonin was performed using the protocol adapted from [105]. The harvested samples were resuspended in PBS (phosphate-buffered saline, pH 7.4) with 250 mM sucrose and 1x PIC (Roche). Protein concentration was estimated using the Bradford method with the day 2 harvested samples, i.e., DMSO-treated and RNAi-induced cells. Following the protein estimation, protein corresponding to ≈25 μg from the DMSO-treated and RNAi-induced samples were treated with increasing amounts of digitonin from 0.05 mg to 1 mg of digitonin/mg protein (diluted in PBS with 250 mM sucrose). Cells treated with 1% Triton-X 100, which solubilizes all membranes, represent complete release of proteins and serve as a positive control. After adding digitonin, the suspension was vortexed vigorously and incubated at 37 °C for 2-3 min with shaking (Thermo mixer-600 rpm). The incubated sample was centrifuged at 20,800 g for 15 min at 4 °C (Rotor F45-30-11, Eppendorf), and the resulting supernatant was analyzed by immunoblotting using various antibodies.

### Immunoblotting

The samples or cell lysate to be analyzed were denatured in Laemmli buffer for 5 min at 95°C and separated by SDS-PAGE. After the gel electrophoresis, immunoblotting was performed as described in [65]. The following primary antibodies were used in this study: mouse α-GFP (Sigma, 1:2,000), mouse α-MtHSP70 (1:1,000); rabbit α-*Tb*Enolase (cytosolic marker, 1:20,000), <u>glycosomal matrix markers</u>: rabbit α-*Tb*Aldolase (1:20,000), rabbit α-*Tb*Hexokinase, (1:10,000), rabbit α-*Tb*GAPDH (1:10,000), rabbit α-*Tb*PFK (1:10,000), rabbit α-*Tb*G3PDH (1:10,000); <u>glycosomal membrane markers</u>: rabbit α-*Tb*GIM5 and rabbit α-*Tb*PEX11 (both 1:2,000); rabbit α-*Tb*VDAC (mitochondrial membrane marker, 1:10,000), rabbit α-*Tb*BiP (ER marker, 1:5,000), rabbit α-*Tb*PEX5 (1:20,000) and rabbit α-Human alpha-tubulin (Abcam, 1:2,000) which shows cross-reactivity with *Trypanosoma* tubulin. Polyclonal antibodies against *Tb*PEX15 were generated in this study. All the above antibodies were prepared in PBS-T buffer (PBS, pH 7.4, 0.05% Tween-20) with 3% BSA except for VDAC, MtHSP70, and alpha-tubulin, which were prepared with 2.5% and 5% fat free milk, respectively. The corresponding secondary antibodies i.e., goat anti-rabbit IRDye 680 or goat anti-mouse IRDye 800CW (LI-COR Biosciences, 1:15,000) were used. Immunoblots were scanned using the LI-COR Odyssey Infrared Imaging System and analyzed with Image Studio version 5.2.

### *Tb*PEX15 antibody production

*Tb*PEX15 ΔTMD_1-320aa_ was subcloned to pET28a vector with TEV cleavage site. The plasmid was transformed to *E. coli* BL21(DE3) pRIL for expression. Protein production was induced with 0.5 mM IPTG at 18°C for 16 h. Cells were harvested and stored at −80°C until purification. For purification, the cells were thawed on ice and resuspended in lysis buffer (50 mM Tris, 150 mM NaCl, pH 7.9, 1 mM DTT, 25 μg/mL DNase I, 50 μg/mL lysozyme, 1 mM PMSF, 5 μg/mL Antipain, 2 μg/mL Aprotinin, 0.35 μg/mL Bestatin, 6 μg/mL Chymostatin, 2.5 μg/mL Leupeptin, 1 μg/mL Pepstatin). Cell lysis was performed by EmulsiFlex. The suspension was centrifuged at 4°C and 4,500 rpm for 15 min (rotor SX4400, Beckman Coulter). The supernatant was transferred to fresh tubes and centrifuged at 4°C 14,000 rpm for 1 h (rotor SS-34, Thermo Scientific). The supernatant was loaded onto an immobilized nickel affinity chromatography column (HisTrap, GE Healthcare). Gradient elution with up to 300 mM imidazole in buffer A (50 mM Tris, 150 mM NaCl, pH 7.9) was performed to retrieve the protein which was further treated with TEV protease for cleavage of the affinity tag at 4°C overnight. Reverse nickel affinity chromatography was performed followed by size exclusion chromatography using buffer A for antiserum production or buffer B (PBS, pH 7.4) for affinity purification. The purified protein was concentrated and snap-frozen in liquid nitrogen and stored at −80°C before it was sent to Eurogentec as antigen for polyclonal antibody production in rabbit. The received antiserum was able to detect the antigen in immunoblots but also detected multiple unspecific bands when tested on BSF 90-13 lysates. Hence, the antiserum was affinity purified by Eurogentec using recombinant *Tb*PEX15 ΔTMD_1-320aa_. The antibody was used at a concentration of 1 μg/mL in TBS-T buffer without addition of BSA or fat free milk.

#### Microscopy

Microscopy images were quantified using Pearson’s correlation coefficient, which was calculated with the Zen 3.6 pro colocalization tool (blue edition). Statistical significances were calculated using a one-way ANOVA (mixed) by Dunnett’s multiple comparisons test, with each row representing matched or repeated measures.

#### Immunoblots

Immunoblot signal intensities were quantified using Empiria Studio 2.3 software. α-Tubulin was used as an internal loading control. One-way ANOVA analysis was performed for statistical analysis using GraphPad Prism 10. Each induced sample was normalized to its respective DMSO control. The significance levels are indicated as follows: ****adjusted p<0.0001, ***adjusted p<0.001, **adjusted p<0.01, *adjusted p<0.05.

## Data availability

The mass spectrometry proteomcis data have been deposited to the ProteomeXchange Consortium [106] via the PRIDE [107] partner repository and are accessible using the dataset identifier PXD057212.

## Supporting information

Description of additional supplementary files

## Acknowledgements

This project work has received funding from the European Union’s Horizon 2020 research and innovation program under the Marie Skłodowska-Curie grant agreement No. 812968 (CK, HD, BW and RE). The work was supported by DFG grant ER178/17-1 to RE. We thank Paul Michels for kindly providing various antibodies, Lavanya Mahadevan for providing the PEX15 and TA2 cell lines, and the PRIDE team for data deposition to the ProteomeXchange Consortium. We would like to acknowledge the use of molecular graphics and analyses performed with UCSF Chimera, developed by the Resource for Biocomputing, Visualization, and Informatics at the University of California, San Francisco, with support from NIH P41-GM103311 and other web tools such as Phyre2, Sequence Identity and Similarity (SIAS) tool.

## Author contributions

CK, HD, BW, VK, RE conceived and planned the experiments. CK, HD, LH performed all experiments. WS, SO, BW, VK, RE supervised the work. CK, HD, SO and VK wrote the manuscript with support of RE and BW. All authors read and gave feedback on the manuscript.

