## Supplementary figures and tables for "High confidence glycosomal membrane protein inventory unveils trypanosomal Peroxin PEX15"

### Supplementary Information File

### Supplementary Figures

a

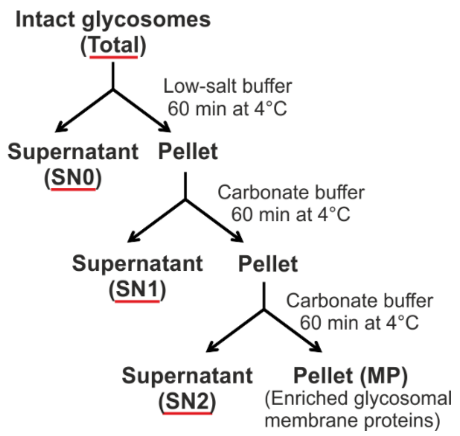

**Fig. S1: Workflow of membrane protein enrichment from the isolated intact glycosomes**. **a** Samples were taken at various steps and analyzed by immunoblotting (**Fig. 1c**) and LC-MS (**Fig. 1d**). Total: Intact glycosomes; SN0: Supernatant post low-salt treatment; SN1/SN2: Supernatant post first or second carbonate treatment; MP: Integral membrane-protein pellet.

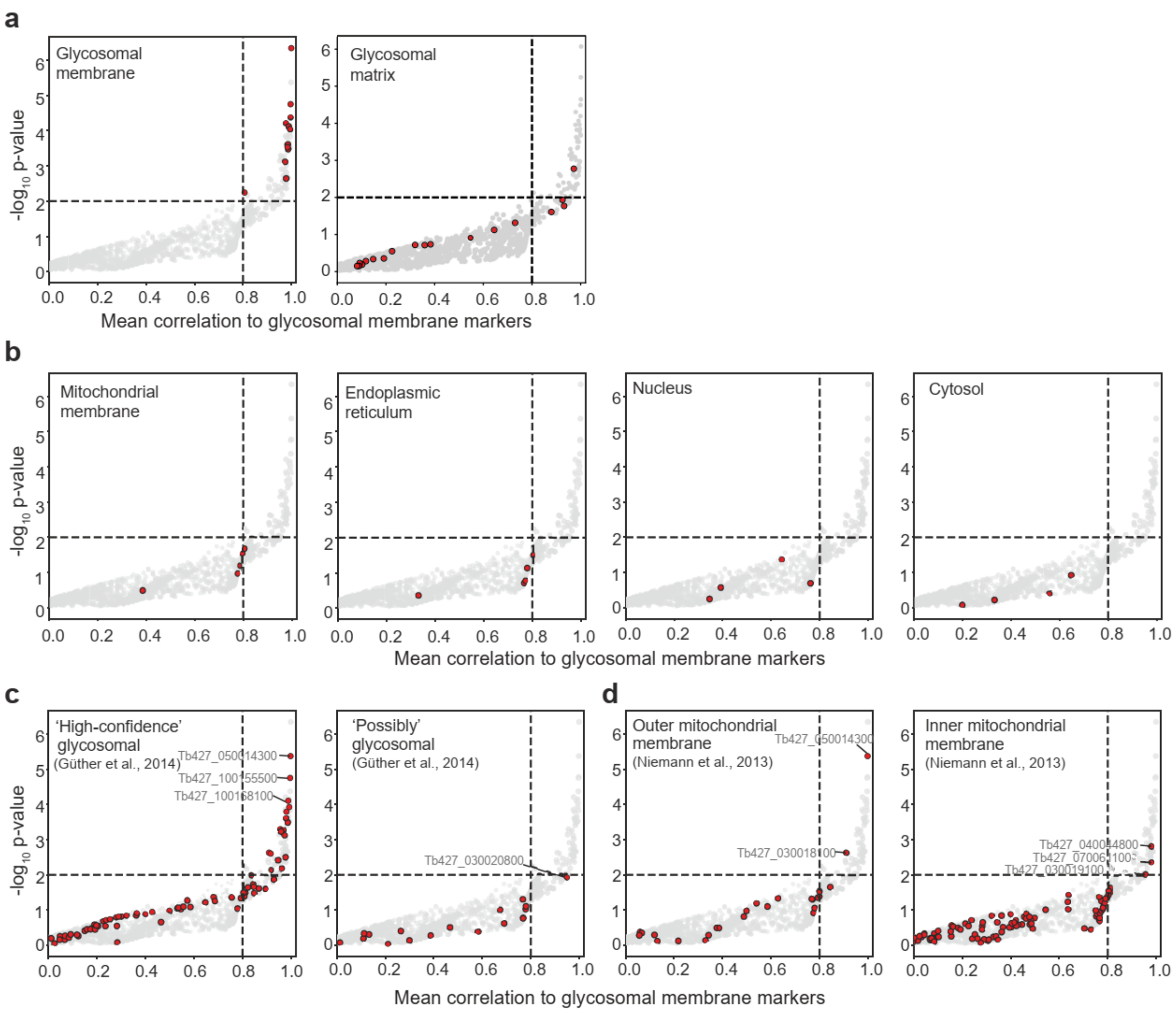

**Fig. S2: Evaluation of the PCP approach used to establish a preliminary list of glycosomal membrane proteins.** Rank product plot as shown in **Fig. 2b** highlighting, **a** glycosomal membrane and matrix marker proteins, **b** marker proteins of different subcellular compartments as indicated, **c** proteins reported to be high-confident glycosomal proteins or "possibly glycosomal" in a previous study of affinity-purified glycosomes (Güther et al. 2014) and **d** proteins present in the outer or inner mitochondrial membrane according to Niemann et al. (2013). See **File S1** for marker proteins used in this study. **a - d,** Horizontal and vertical dashed lines indicate a p-value of 0.01 and a mean correlation of 80%, respectively.

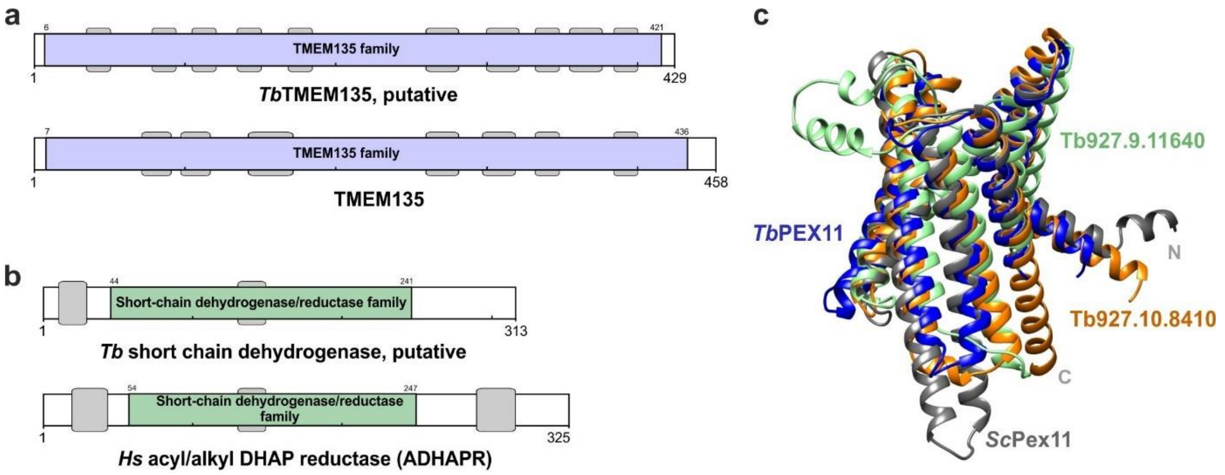

**Fig. S3: a** Schematic representation of the domain architecture of the putative *Tb*TMEM135 (*T. brucei*) and *Hs*TMEM135 (*H. sapiens*). The TMEM135 family domain is well conserved and shown in blue. **b** Schematic representation of the putative *Tb* short-chain dehydrogenase (SDR) domain architecture and *Hs* acyl/alkyl DHAP reductase (ADHAPR). The SDR family domain is well conserved and is shown in green. The scheme represents the transmembrane segments and domain architecture based on the InterPro Scan. **c** Structural comparison of the putative *Tb*PEX11 family protein (orange) with PEX11 from *T. brucei* (blue) and *S. cerevisiae* (gray), using the *Sc*PEX11 structure as reference.

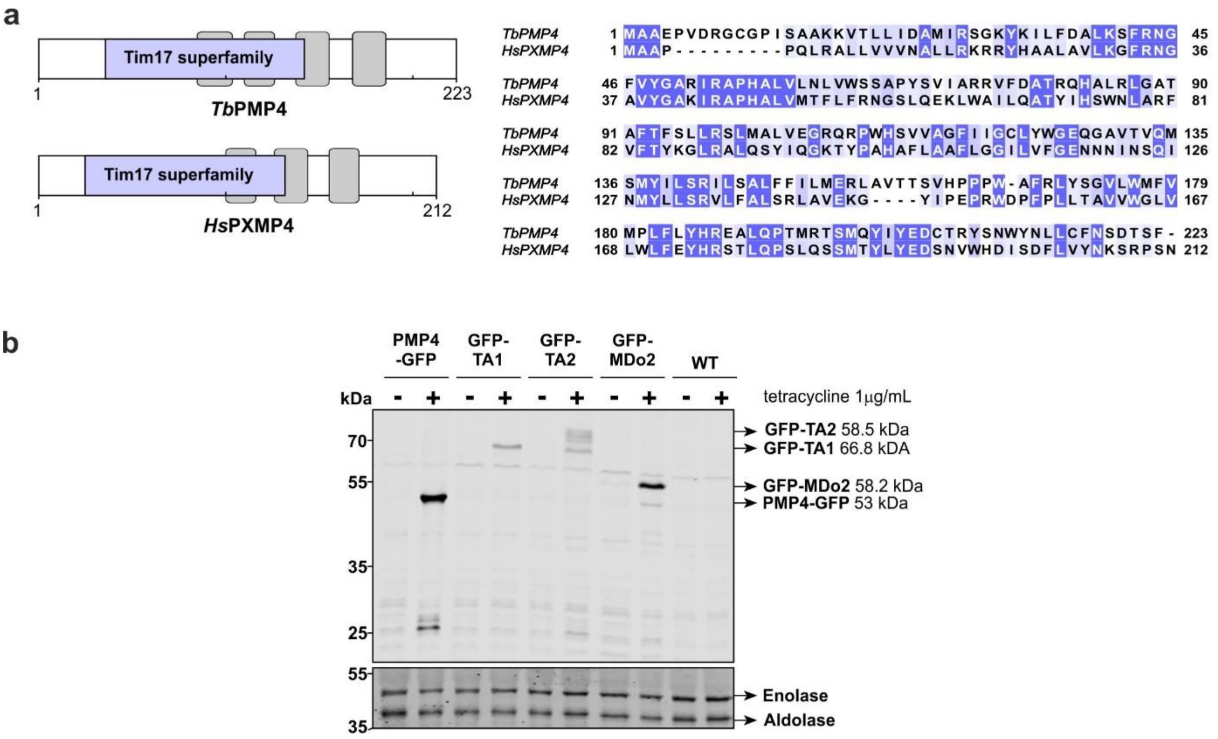

**Fig. S4: Domain architecture of the validated glycosomal membrane proteins. a** Left panel: Schematic representation of the domain architecture of the putative *Tb*PMP4 (*T. brucei*) and *Hs*PXMP4 (*H. sapiens*). The scheme represents the transmembrane segments predicted using Phobius webtool, and domain architecture based on the InterPro Scan. The Tim17 superfamily domain is well conserved and is shown in blue. Right panel: Multiple sequence alignment of *Tb*PMP4 (TriTrypDB ID Tb427_090008200) and *Hs*PXMP4 (UniProt ID Q9Y6I8) protein sequence. The sequence conservation is colored according to the percentage identity with conservation threshold of 30%. **b** Expression of tetracycline-inducible N/C-terminally GFP-tagged PMPs in procyclic form (PCF) trypanosomes (shown in **Fig. 3c**) was verified by immunoblotting using monoclonal antibodies against the GFP tag (upper panel). Enolase and aldolase served as loading controls (lower panel).

a

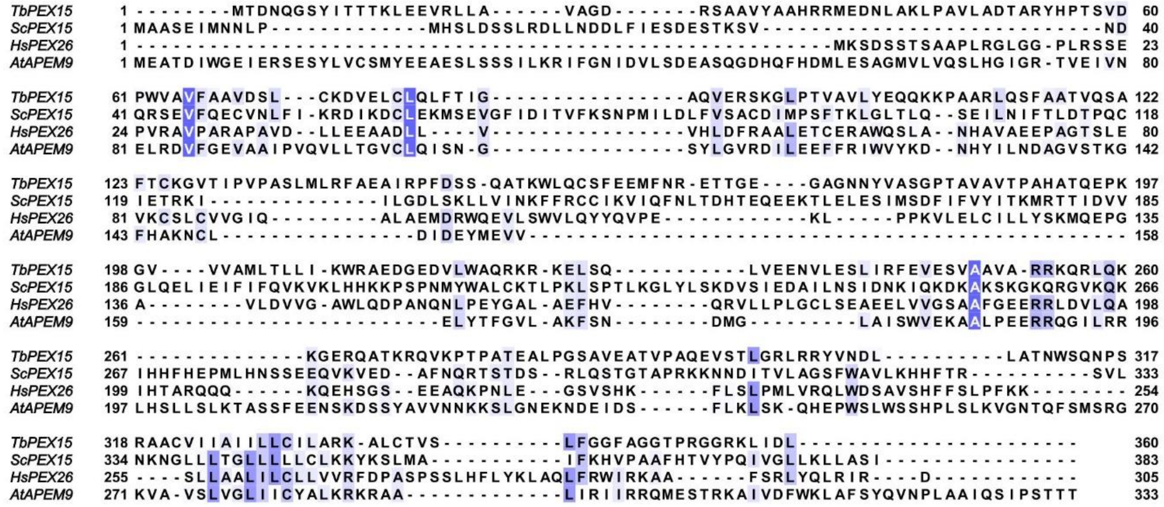

b

| | *T. brucei* | *S. cerevisiae* | *H. sapiens* | *A. thaliana* | |
|---|---|---|---|---|---|
| *T. brucei* | | 23.05 | 25.9 | 22.82 | % Similiarity |
| *S. cerevisiae* | 11.94 | | 22.62 | 25.52 | |
| *H. sapiens* | 16.39 | 11.14 | | 27.54 | |
| *A. thaliana* | 13.21 | 12.91 | 18.68 | | |
| | % Identity | | | | |

c

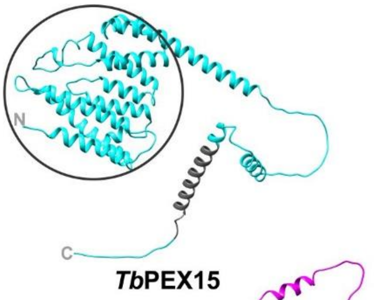

d

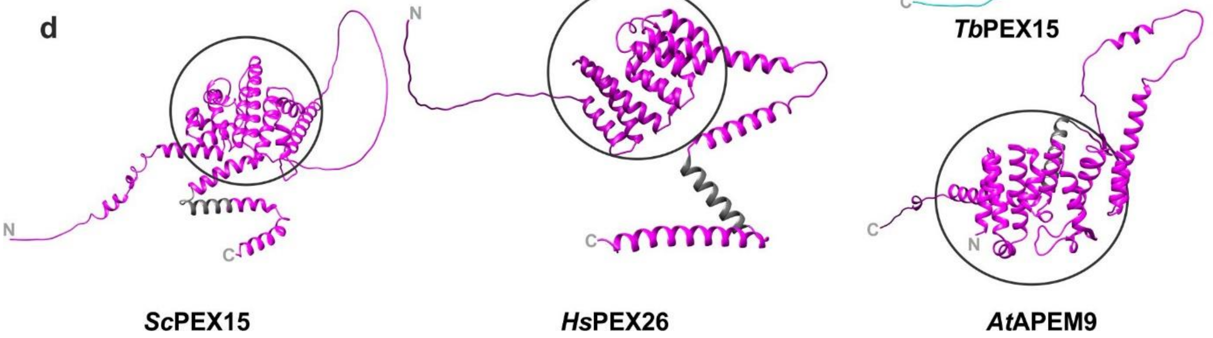

**Fig. S5: Bioinformatic analysis of PEX15/PEX26 orthologs across organisms. a** Multiple sequence alignment of *Tb*PEX15 (*T. brucei*), *Sc*PEX15 (*S. cerevisiae*), *Hs*PEX26 (*H. sapiens*) and *At*APEM9 (*Arabidopsis thaliana*) protein sequences. *Tb*PEX15 protein sequence was obtained from TriTrypDB with accession ID Tb427_100021800 whereas yeast, human and plant protein sequences were obtained from UniProt with UniProt ID Q08215 (*S. cerevisiae*), Q7Z412 (*H. sapiens*), Q8W4B2 (*Arabidopsis thaliana*). Multiple Sequence alignment was performed using the Clustal Omega tool, and the aligned sequences were visualized using JalView software. The sequence conservation is colored according to the percentage identity with conservation threshold of 30%.. The conserved C-terminal domain comprises the predominant portion of the TMD across organisms (underlined). **b** Percentage identity and similarity matrix of PEX15 counterpart across organism based on SIAS homology modelling (http://imed.med.ucm.es/Tools/sias.html). **c,d** AlphaFold predicted structures of PEX15 (*T. brucei* and *S. cerevisiae*), PEX26 (*H. sapiens*), and APEM9 (*Arabidopsis thaliana*) show high similarity. The black circles represent the common helical bundle, which is known to interact with PEX6. The predicted structures are presented with a rainbow color scheme, where the N-terminus is blue, and the C-terminus is in red color. The characteristic predicted C-terminal TMD and the putative hydrophobic membrane-spanning segment of *Hs*PEX26 (citation) are marked in grey.

**a**

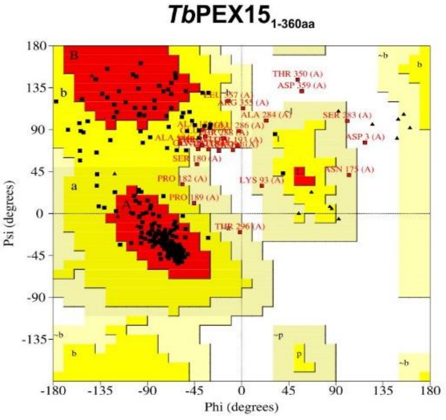

**b**

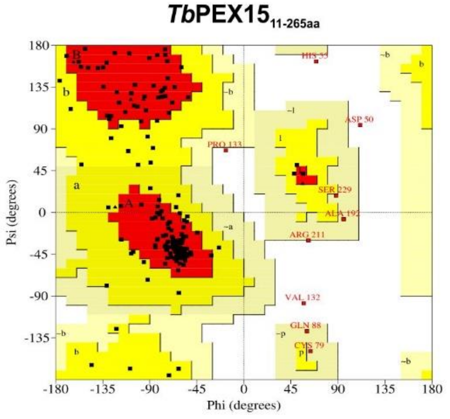

Plot statistics

| | | |
|---|---|---|
| Residues in most favoured regions [A,B,L] | 258 | 80.4% |
| Residues in additional allowed regions [a,b,l,p] | 40 | 12.5% |
| Residues in generously allowed regions [~a,~b,~l,~p] | 17 | 5.3% |
| Residues in disallowed regions | 6 | 1.9% |
|  | ---- | ------ |
| Number of non-glycine and non-proline residues | 321 | 100.0% |
| Number of end-residues (excl. Gly and Pro) | 2 | |
| Number of glycine residues (shown as triangles) | 20 | |
| Number of proline residues | 17 | |
|  | ---- |  |
| Total number of residues | 360 | |

Plot statistics

| | | |
|---|---|---|
| Residues in most favoured regions [A,B,L] | 164 | 80.8% |
| Residues in additional allowed regions [a,b,l,p] | 31 | 15.3% |
| Residues in generously allowed regions [~a,~b,~l,~p] | 4 | 2.0% |
| Residues in disallowed regions | 4 | 2.0% |
|  | ---- | ------ |
| Number of non-glycine and non-proline residues | 203 | 100.0% |
| Number of end-residues (excl. Gly and Pro) | 2 | |
| Number of glycine residues (shown as triangles) | 7 | |
| Number of proline residues | 9 | |
|  | ---- |  |
| Total number of residues | 221 | |

**Fig. S6: Validation of modelled structures using Ramachandran plot. a** The Ramachandran plot for *Tb*PEX15 protein (full-length) structure modelled using AlphaFold2. **b** Likewise, the Ramachandran plot for *Tb*PEX15 protein (11-265aa), modelled using the Phyre2 tool with the crystal structure of *Sc*PEX15 (43-259aa) as a template. In both plots, ≤ 2% of residues were observed in the disallowed region (white area) of the Ramachandran plot. Ramachandran plots were generated using the PROCHECK tool from the Saves server (https://saves.mbi.ucla.edu/).

a

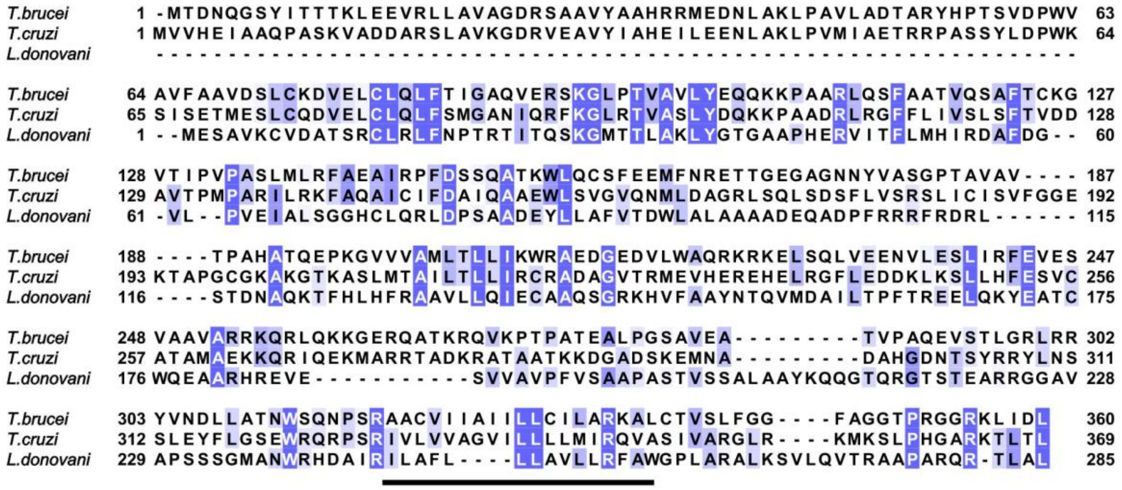

b

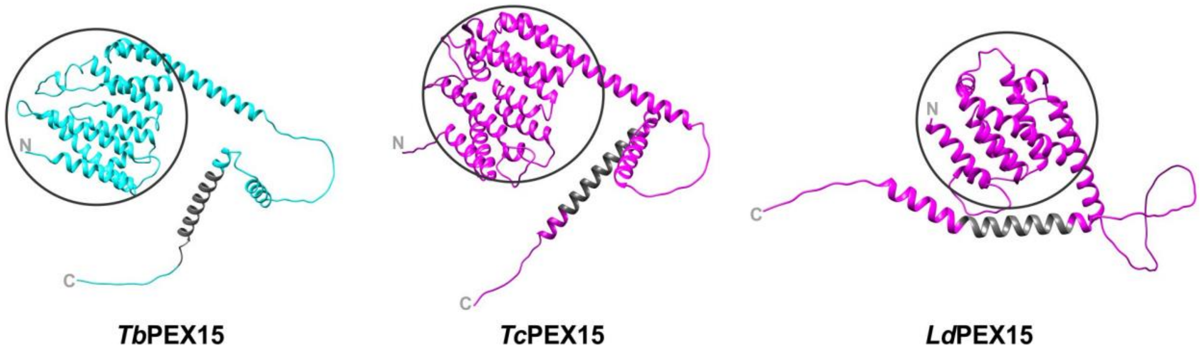

**Fig. S7: Bioinformatic analysis of PEX15 orthologs of parasites. a** Multiple sequence alignment of *Tb*PEX15 (*T. brucei*), *Tc*PEX15 (*T. cruzi*), and *Ld*PEX15 (*L. donovani*) protein sequences. Parasite protein sequences were obtained from TriTrypDB with accession IDs Tb427_100021800 (*T. brucei*), TcCLB.508059.20 (*T. cruzi*), and LdBPK_210110.1 (*L. donovani*). Multiple sequence alignment was performed using the Clustal Omega tool, and the aligned sequences were visualized using JalView software. The sequence conservation is colored according to the percentage identity with conservation threshold of 30%. Alignment indicates that *Trypanosoma* (*Tb* and *Tc*) PEX15 have longer N-terminal extension than the *Leishmania* counterparts. The conserved C-terminal domain comprises the predominant portion of the predicted TMD (underlined). **b** AlphaFold predicted structures of PEX15 (*T. brucei*, *T. cruzi*, and *L. donovani*) show high similarity. The black circles represent the common helical bundle similar to yeast and human counterpart. The predicted structures are presented with a rainbow color scheme, where the N-terminus is blue, and the C-terminus is red in color. The predicted C-terminal TMD is marked in grey.

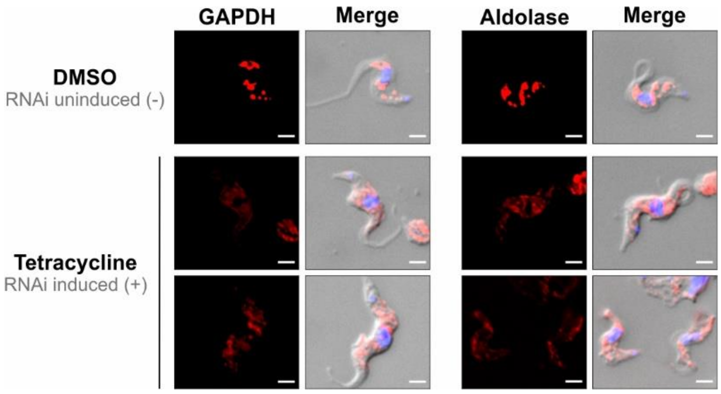

**Fig. S8: Immunofluorescence microscopy analysis of glycosomes upon PEX15 RNAi.** On day 2 of PEX15 RNAi, both DMSO and tetracycline treated cells were analyzed for aldolase and GAPDH by immunofluorescence microscopy. In DMSO-treated cells, aldolase and GAPDH display a punctate glycosomal pattern. Upon PEX15 RNAi induction, glycosomal markers labelling was similar but puncta appeared less bright and more diffuse. Merge channel additionally includes DAPI (DNA stain) and brightfield. The scale bar represents 2 μm.

### Supplementary Tables

**Table S1: Strains and Plasmids**

| Sl no. | Expression in | Construct | Primer pair | Restriction sites | Cloned in vector |
|---|---|---|---|---|---|
| 1 | *S. cerevisiae* | GAL4 BD-*Tb*PEX15$_{1\text{-}360aa}$ | RE8663 – RE8664 | SalI/NotI | pPC97 |
| 2 | *S. cerevisiae* | GAL4 BD-*Tb*PEX15ΔTMD$_{1\text{-}320aa}$ | RE8663 – RE8665 | SalI/NotI | pPC97 |
| 3 | *S. cerevisiae* | GAL4 BD-*Tb*PEX15$_{1\text{-}360aa}$ (LL 19,20 SS) | RE8755 – RE8756 | Quick change PCR | pPC97 |
| 4 | *S. cerevisiae* | GAL4 BD-*Tb*PEX15ΔTMD$_{1\text{-}320aa}$ (LL 19,20 SS) | RE8755 – RE8756 | Quick change PCR | pPC97 |
| 5 | *S. cerevisiae* | GAL4 AD-*Sc*PEX5 | (Kerssen *et al.* 2006) | | |
| 6 | *S. cerevisiae* | GAL4 BD-*Sc*PCS60 | (Effelsberg *et al.* 2016) | | |
| 7 | *S. cerevisiae* | GAL4 AD-*Tb*PEX6 | (Mahadevan *et al.* 2024) | | |
| 8 | *T. brucei* | *Tb*PMP4-GFP | RE7839 – RE7840 | HindIII/BamHI | pGN1 |
| 9 | *T. brucei* | GFP-*Tb*PEX15 (TA1) | RE8896 – RE8897 (vector) / RE8898 – RE8899 (insert) | In-fusion cloning | pGC1 |
| 10 | *T. brucei* | GFP-*Tb*TA2 | RE7476 – RE7477 | HindIII/BamHI | pGC1 |
| 11 | *T. brucei* | GFP-*Tb*MDo2 | RE7579 – RE7578 | HindIII/BamHI | pGC1 |
| 12 | *T. brucei* | GFP-PTS1 | (Dawidowski *et al.* 2017) | | |
| 13 | *T. brucei* | *Tb*PEX15 RNAi (Fragment 1) | RE8659 – RE8660 | BamHI/MfeI | Ligate fragments 1 & 2 |
| 14 | *T. brucei* | *Tb*PEX15 RNAi (Fragment 2) | RE8661 – RE8662 | HindIII/MfeI | |
| 15 | *T. brucei* | *Tb*PEX15 RNAi (Stem-loop) | Ligated fragments 1 & 2 | BamHI/HindIII | p2T7-177 |
| 16 | *E. coli* | His$_6$-*Tb*PEX15 ΔTMD$_{1\text{-}320aa}$ | RE8452 – RE8453 (vector) / RE8666 – RE8948 (insert) | In-fusion cloning | pET28a |

**Table S2: Oligonucleotides**

| Primer | Sequence 5' to 3' |
|---|---|
| RE7578 | AGAGGATCCCTACAATTTGGAGGGCTGCGTGCAGT |
| RE7579 | AAGAAAGCTTCCATGGTGTCGTACGCCTCTGCATT |
| RE7839 | AAGACAAGCTTATGGCGGCTGAGCCCGTGGACAGG |
| RE7840 | GACGGATCCCCAAAGGAAGTATCACTGTTAAAGCA |
| RE8452 | ATGGCTAGCATGACTGGTGGAC |
| RE8453 | ATGGCCCTGAAAATACAGGTTTTCGC |
| RE8659 | AAGACGGATCCATTACGTAGCATCTGGGCCG |
| RE8660 | AAGACCAATTGCAGAGATCGATCAGCTTC |
| RE8661 | AAGACAAGCTTATTACGTAGCATCTGGGCCG |
| RE8662 | AAGACCAATTGACCGTACACAGGGCTTTACG |
| RE8663 | AAGACGTCGACGATGACAGATAATCAAGGTAGCTAT |
| RE8664 | AACAGCGGCCGCTCAGAGATCGATCAGCTTCCT |
| RE8665 | AAGACGCGGCCGCTTAGGCTGCACGAGATGGGTTCTG |
| RE8666 | ACCTGTATTTTCAGGGCCATATGACAGATAATCAAGGTAGCTATATCACTAC |
| RE8755 | GTTCGCAGTTCTGCCGTCGCAGGGGATCG |
| RE8756 | GGCAGAACTGCGAACTTCTTCAAGTTTCGTGG |
| RE8896 | GGATCCACCGGATCTAGATAACTGAT |
| RE8897 | TCGAGATCTGAGTCCGGACTTGTAC |
| RE8898 | TCCGGACTCAGATCTCGAATGACAGATAATCAAGGTAGCTATATCACTACCAC |
| RE8899 | TCTAGATCCGGTGGATCCTCAGAGATCGATCAGCTTCCTTCCG |
| RE8948 | CCACCAGTCATGCTAGCCATTCAGGCTGCACGAGATGGG |
