## Supplementary material for "High confidence glycosomal membrane protein inventory unveils trypanosomal Peroxin PEX15": Description of additional supplementary files

File name: **File S1**

Description: The quantitative MS analysis file provides detailed information on the proteins identified in glycosomal membrane protein enrichment experiments and through the analysis of carbonate-resistant pellets from density gradient fractions

File name: **File S2**

Description: This file presents profiling data for preliminary candidate proteins from Class 1 and Class 2. The analysis encompasses the distributions and abundance profiles of each protein across a density gradient.
