## Supplementary figures and images for "High confidence glycosomal membrane protein inventory unveils trypanosomal Peroxin PEX15"

### File S2

#### Protein gradient profiles

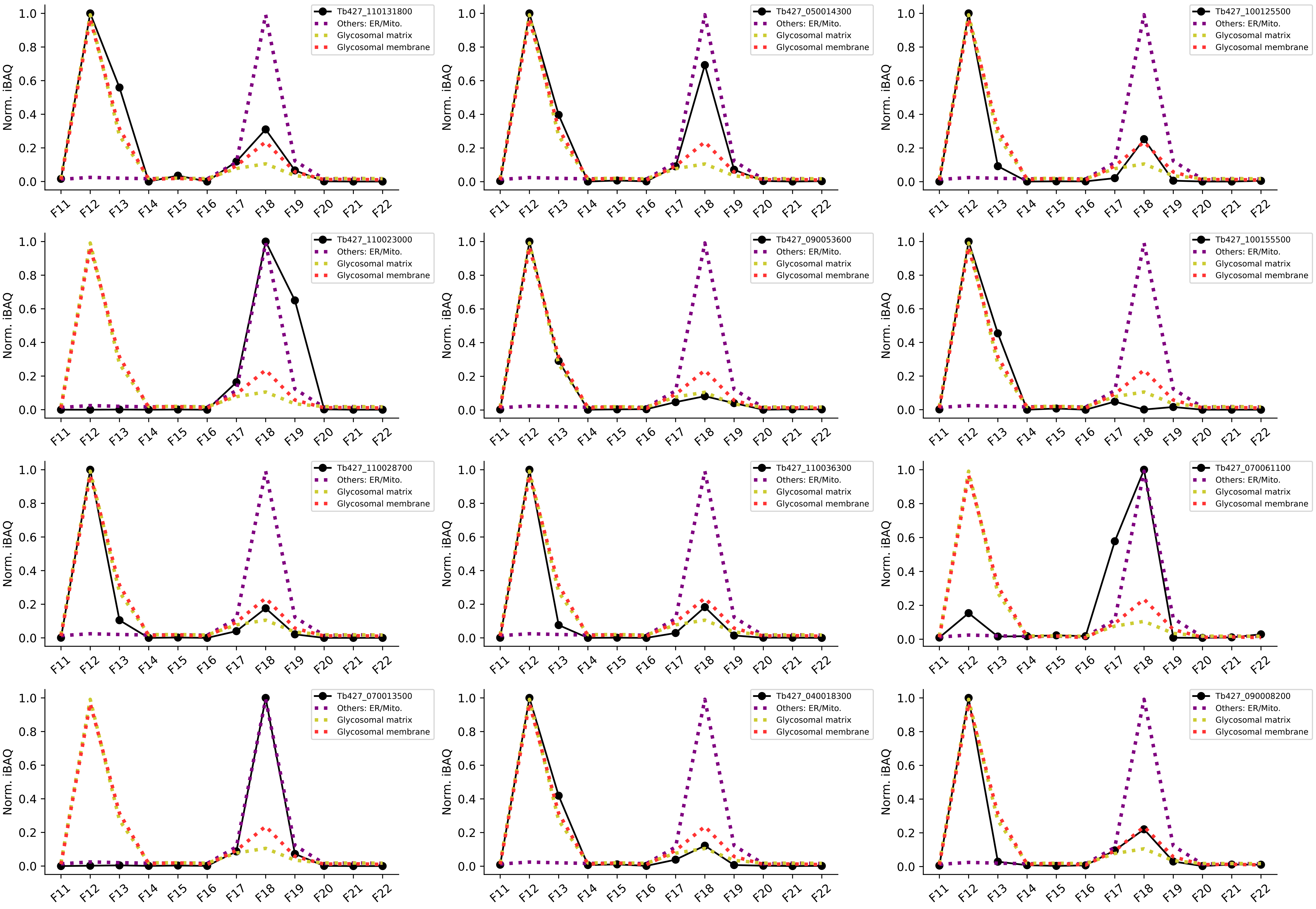

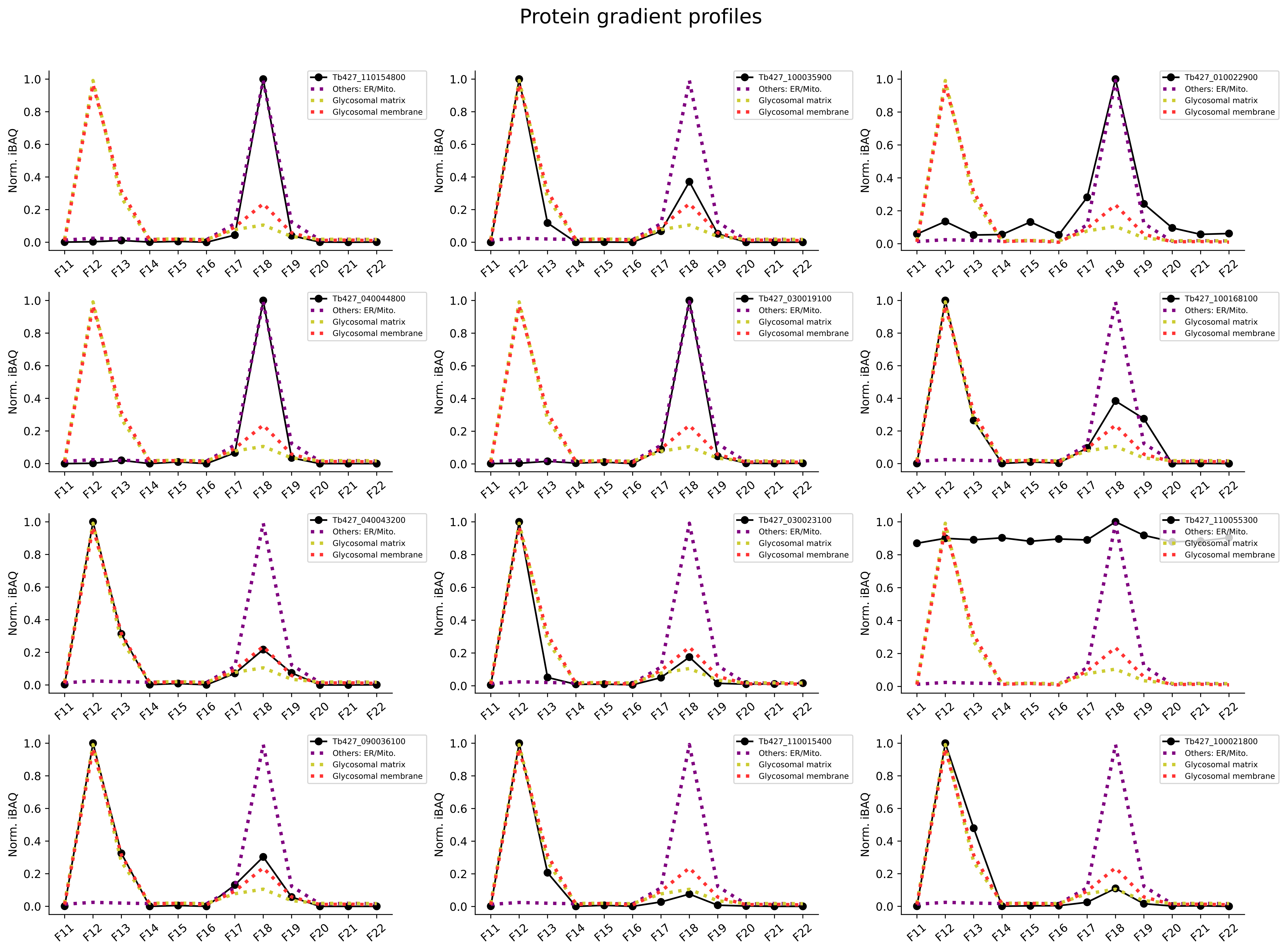

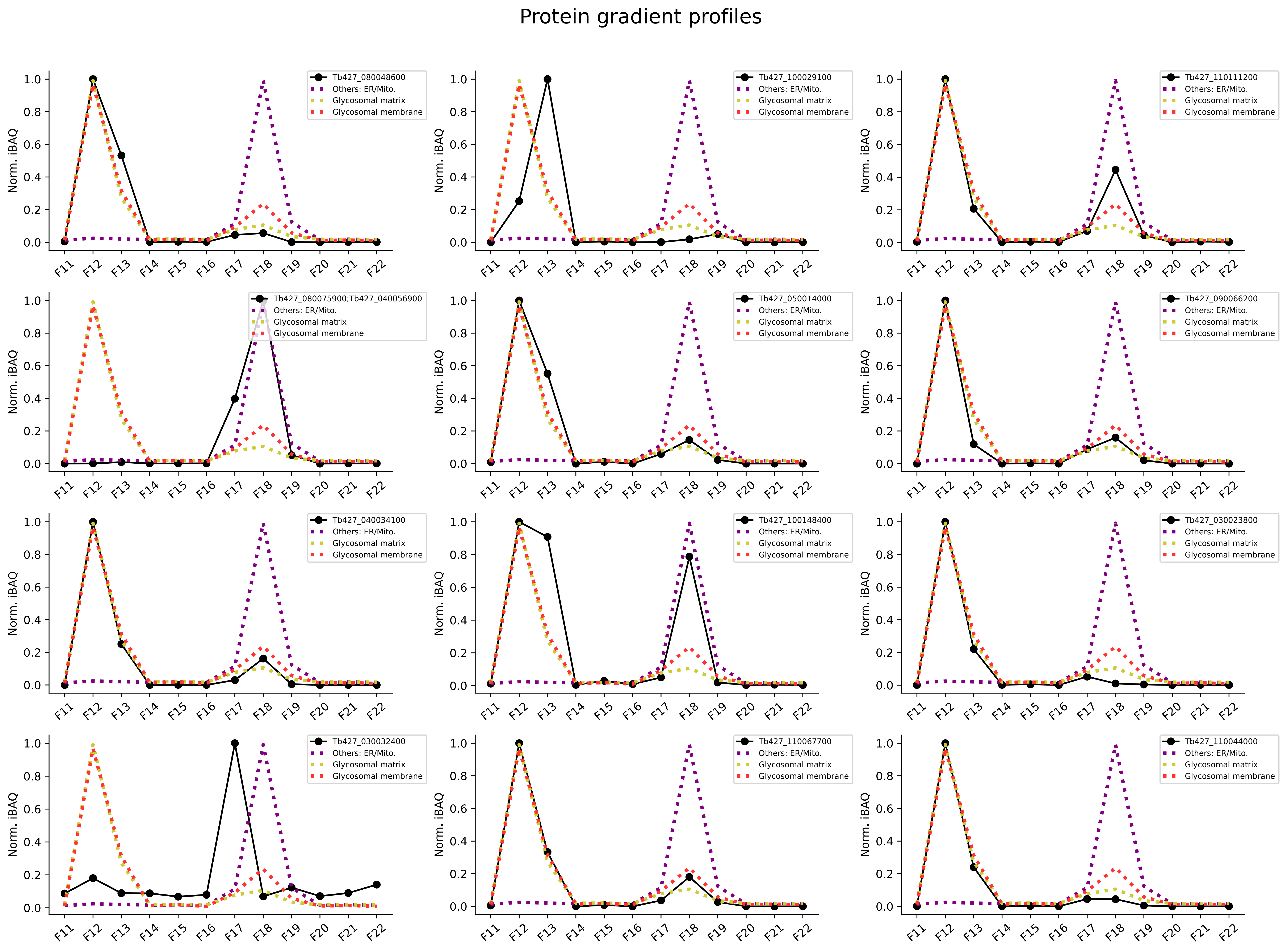

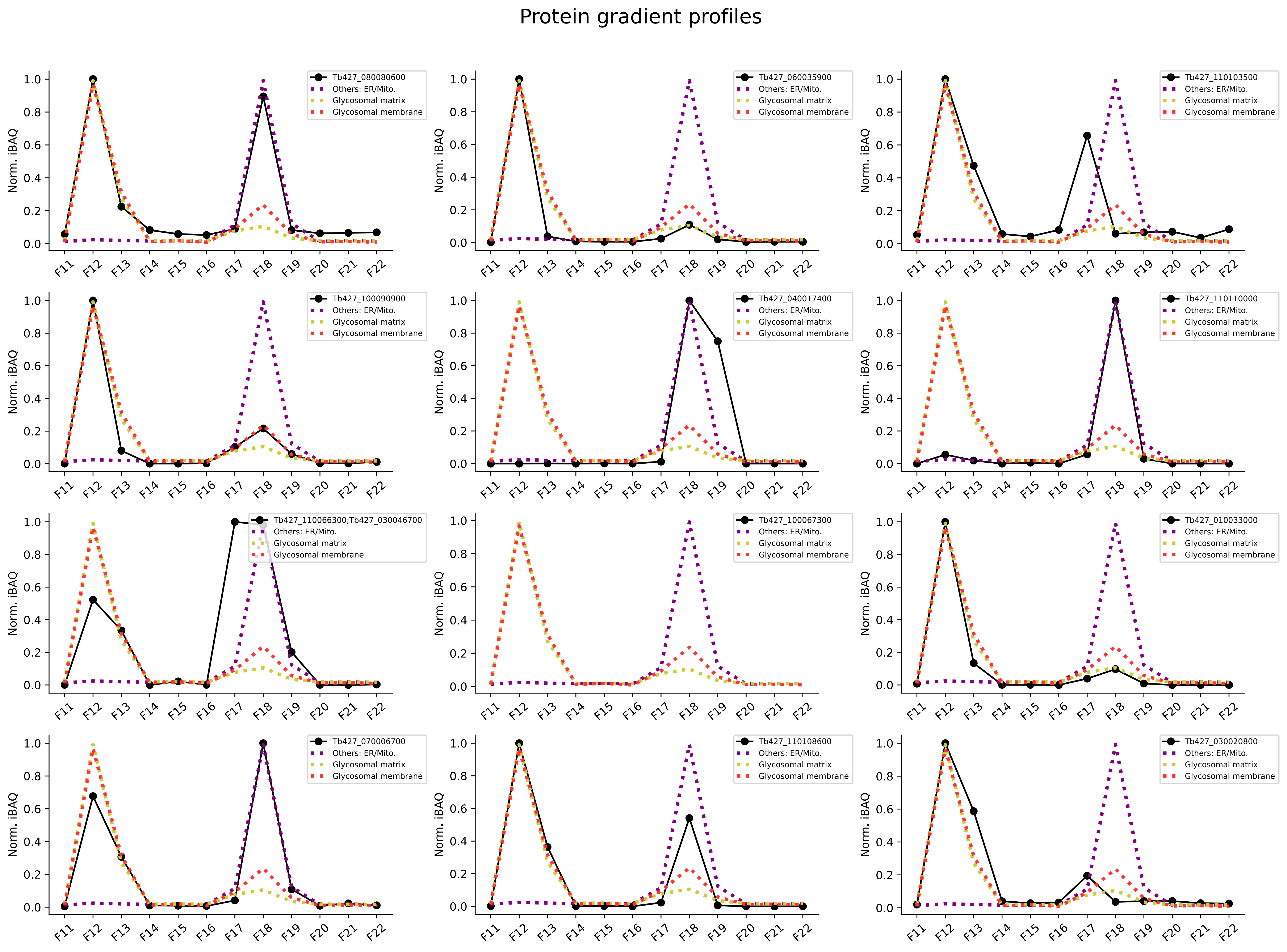

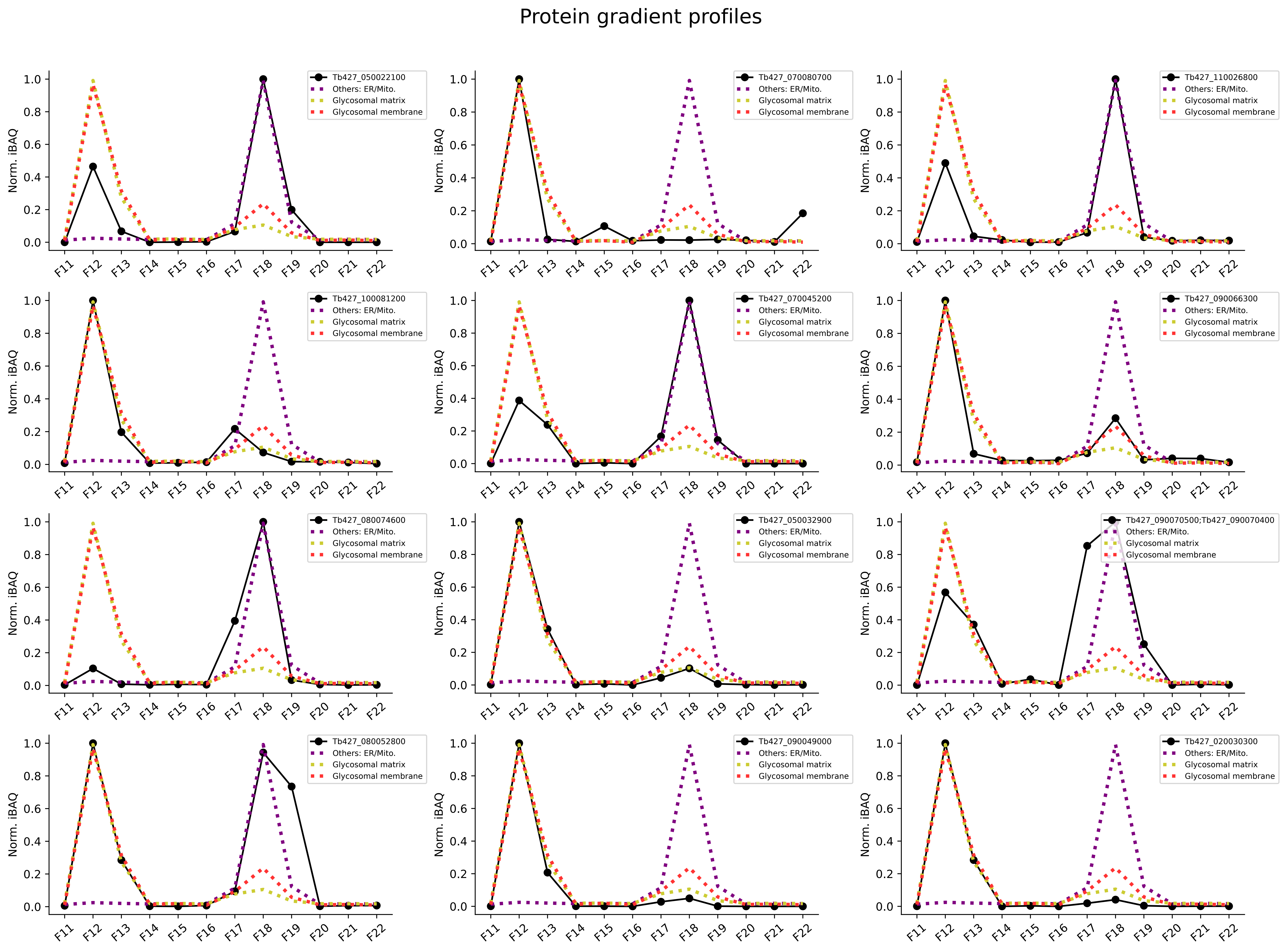

Protein gradient profiles

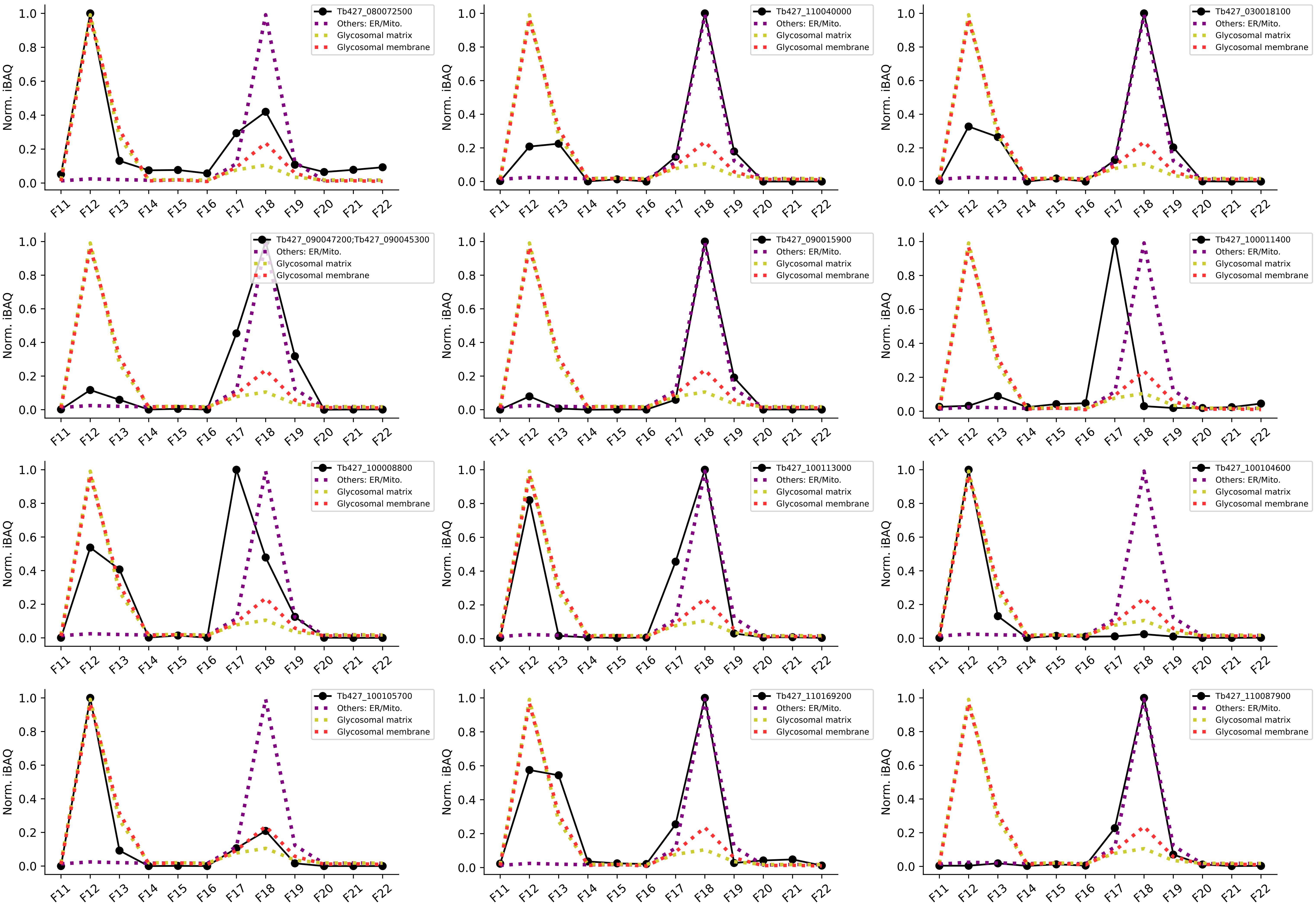

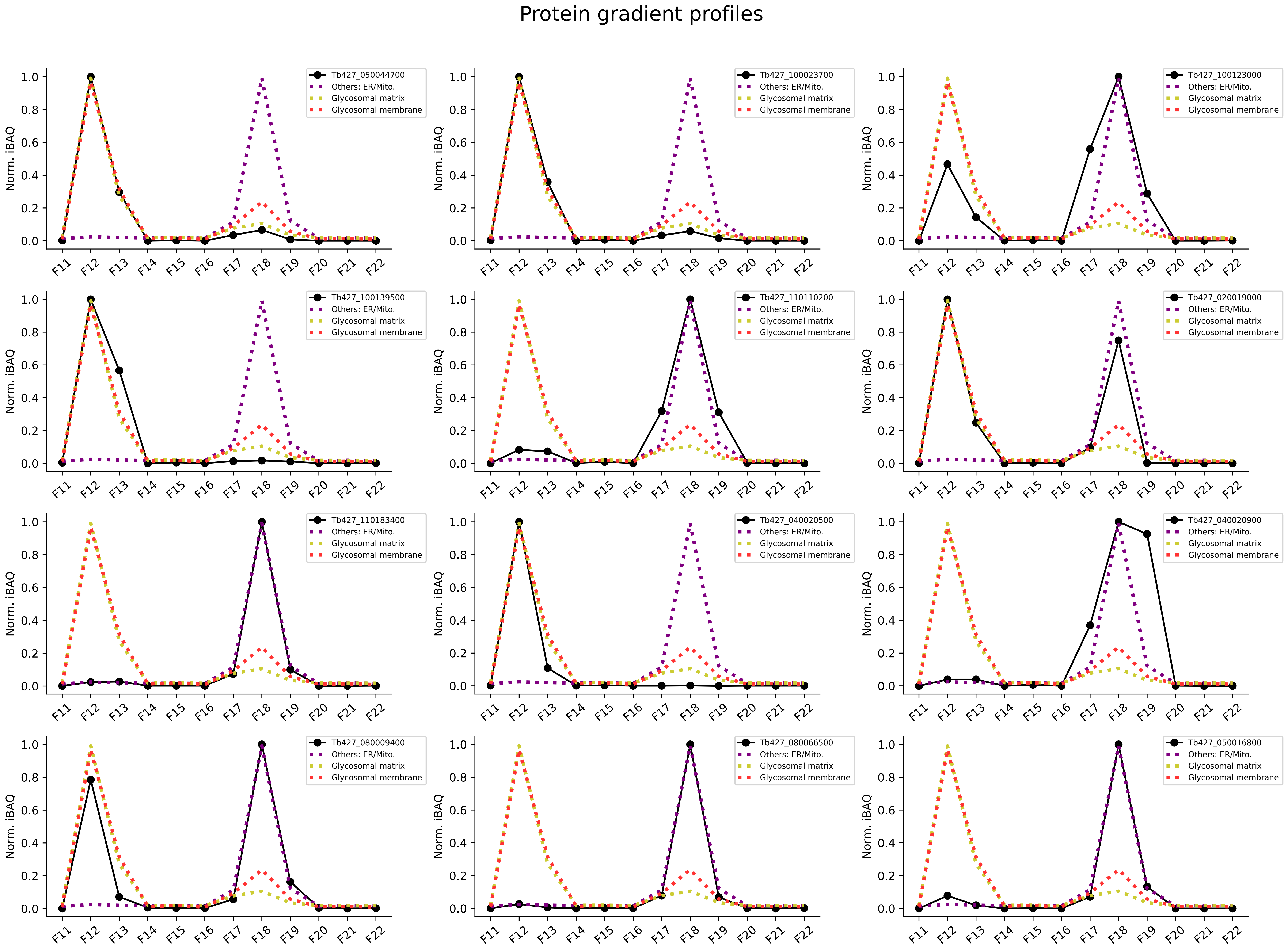

Protein gradient profiles

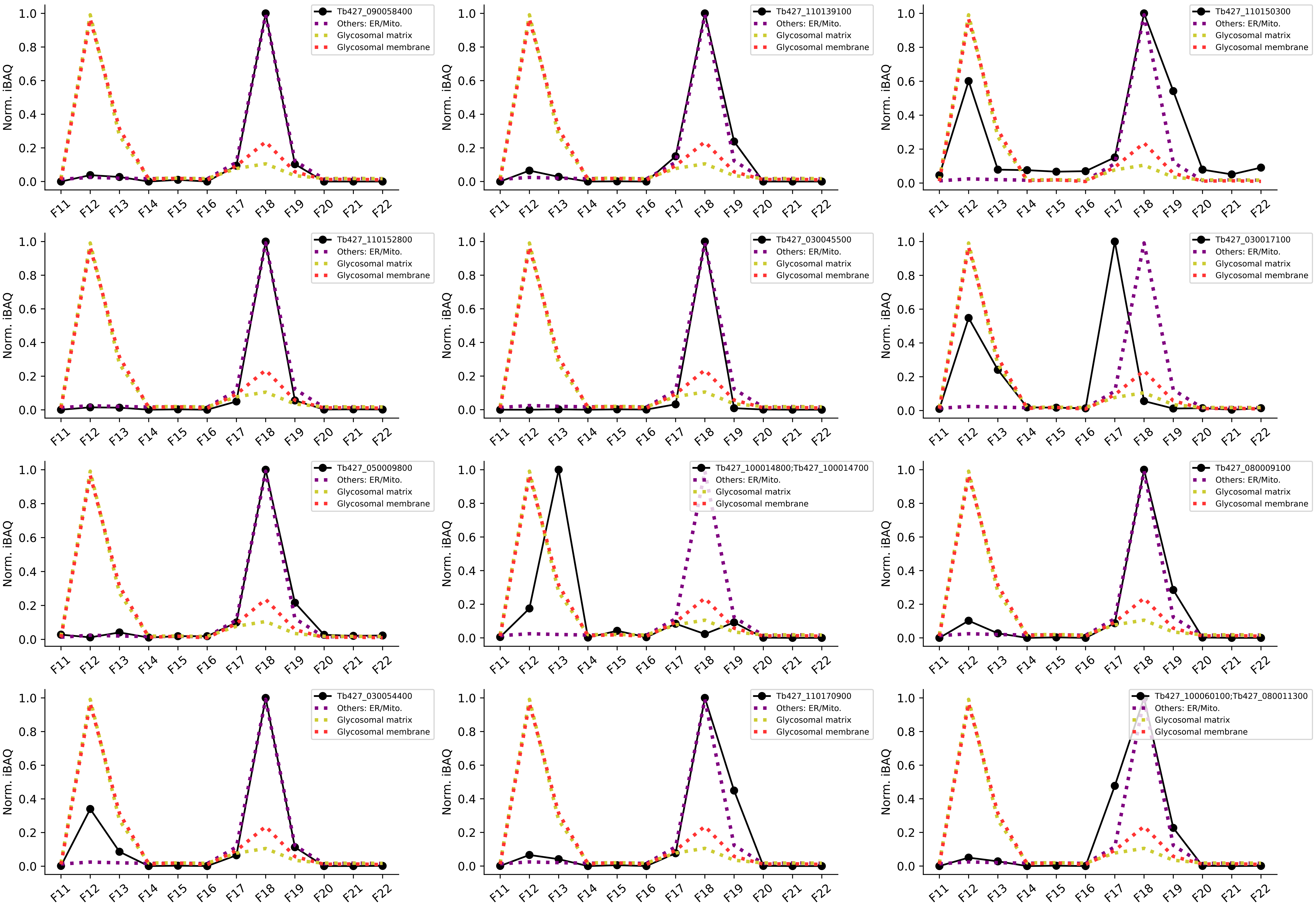

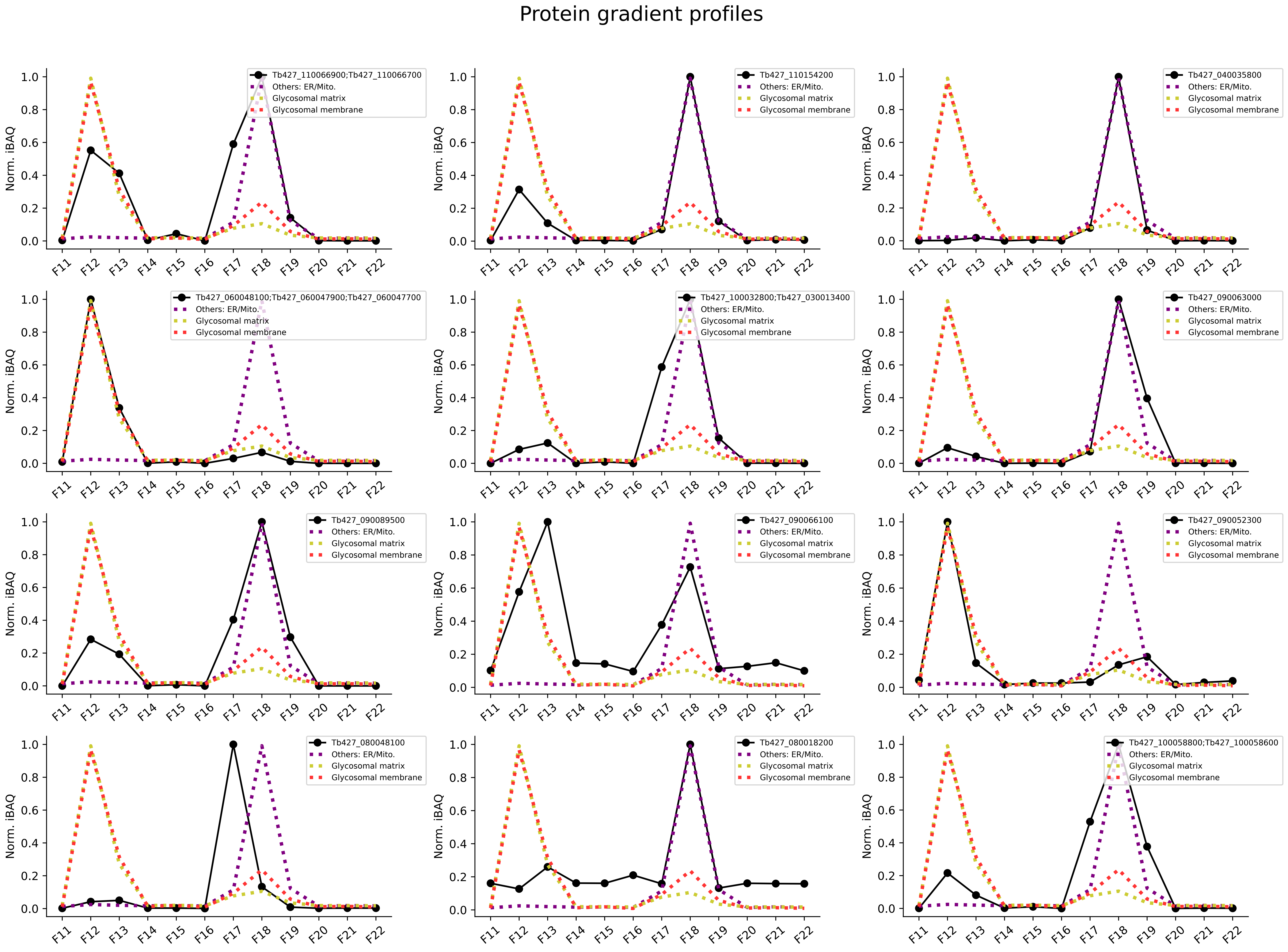

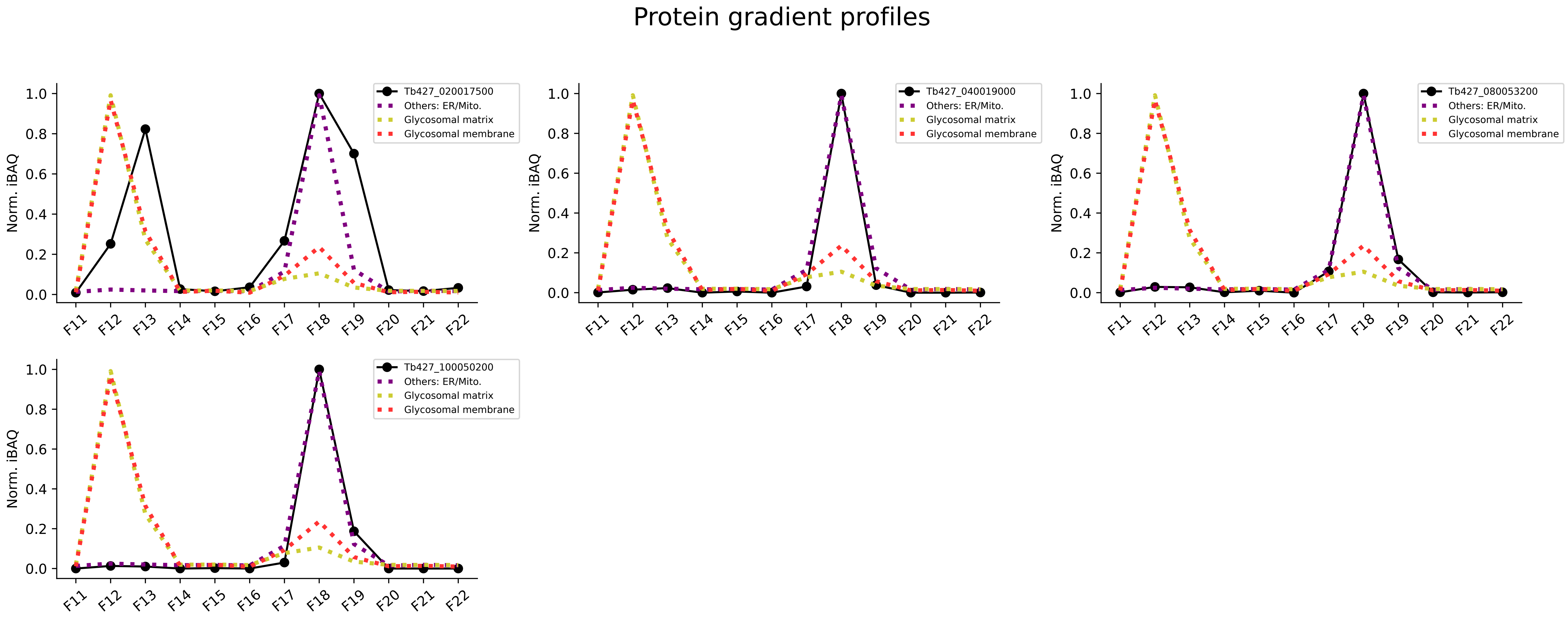
